# Methylglyoxal-derived glycation tunes nuclear viscoelasticity

**DOI:** 10.64898/2026.09.28.754974

**Authors:** Mar Alcaraz-Hurtado, Alysson Duchalet, Angelica Salinas-Torres, Enkela Musliu, Gabriel Jordanov, Jorge Rodriguez-Ramos, Roberto Silva-Rojas, Natacha Fourny, Oscar M. Bautista-Aguilera, Miquel Adrover, Jorge Alegre Cebollada, Luc Bertrand, Andra C. Dumitru

## Abstract

Reactive carbonyl metabolites such as methylglyoxal (MGO) accumulate in diabetes, aging, and metabolic disease, conditions frequently associated with pathological stiffening of human cells and tissues. While glycation of extracellular matrices and cytoskeletal proteins is known to contribute to these mechanical changes, whether the nucleus itself is a direct mechanical target of carbonyl stress remains unknown. Here, by combining advanced atomic force microscopy-based nanomechanical probing with correlative confocal imaging, we show that MGO-derived glycation directly remodels isolated cell nuclei, increasing stiffness, reducing fluidity, and restricting intranuclear deformation propagation. MGO-derived adducts accumulate prominently at the nuclear envelope, and perturbation of lamin A/C and chromatin state reveals that both structures contribute to glycation-induced stiffening. In cells, chronic MGO exposure induces nucleus-intrinsic stiffening and biases YAP nuclear localization in a substrate-stiffness-dependent manner, linking nuclear glycation to altered mechanosensitive regulation. *In vivo*, cardiac nuclear stiffening is only observed along with increased immunodetected MGO-adduct burden, while nuclear fluidity remains largely preserved. Together, these findings establish nonenzymatic glycation as a metabolic mechanism that remodels nuclear mechanics and alters cellular mechanosensing, revealing a previously unrecognized physical consequence of carbonyl stress.

## INTRODUCTION

Pathological alterations in cell and tissue mechanics are central drivers of disease progression in diabetes, aging, and cardiovascular disorders (*1, 2*). Changes in how cells sense, transmit, or dissipate physical forces impair fundamental processes ranging from cell migration and contractility to repair and gene expression (*3*). In the cardiovascular system, chronic hyperglycemia and metabolic dysfunction promote a progressive loss of myocardial compliance that contributes to diastolic dysfunction, yet the molecular events that connect metabolic stress to mechanical stiffening remain incompletely understood (*4–8*). A major biochemical feature of this mechanical phenotype is carbonyl stress, driven by the accumulation of reactive metabolites such as methylglyoxal (MGO) (*9*). Formed primarily as a byproduct of glycolysis and, to a lesser extent, protein catabolism, MGO is a highly reactive dicarbonyl. In type 2 diabetes mellitus, downregulation of its main detoxification pathway, the glyoxalase system, promotes intracellular MGO accumulation and increases carbonyl stress. As a result, MGO reacts extensively and non-enzymatically with nucleophilic residues in proteins and other biomolecules, leading to the formation of advanced glycation end-products (AGEs) that alter protein charge, conformation, function, and supramolecular organization (*10–13*). While AGE-mediated crosslinking of extracellular matrices and intracellular contractile elements is an established contributor to tissue stiffening, whether carbonyl stress directly alters the mechanics of intracellular organelles remains largely unexplored.

The nucleus is a central mechanical organelle. Coupled to the cytoskeleton through the LINC complex, it deforms under physical load, transmits mechanical stress to chromatin and contributes to force-dependent transcriptional regulation (*14, 15*). If carbonyl stress directly alters nuclear material properties, it could therefore affect not only nuclear mechanics, but also the way mechanical cues are converted into transcriptional responses. Nuclear material properties are mainly governed by two interconnected structural compartments: the nuclear lamina and the chromatin (*16*). Both compartments are chemically tunable. Enzymatic post-translational modifications, including lamin A/C phosphorylation and histone acetylation or methylation, remodel nuclear architecture and regulate nuclear mechanical properties and transcriptional accessibility. These examples establish chemical modification as a regulator of nuclear mechanics, but most work has focused on coordinated, enzyme-driven post-translational modifications (PTMs) rather than uncontrolled metabolic chemistry.

Emerging evidence shows that non-enzymatic glycation provides an unregulated chemical input capable of disrupting nuclear mechanics. Many nuclear structural proteins are exceptionally long-lived and enriched in reactive lysine and arginine residues, which makes them highly susceptible to MGO-derived adduct formation. Indeed, recent proteomic and liquid chromatography-tandem mass spectrometry studies have mapped carbonyl-derived AGEs onto basic nuclear proteins, including core histones and chromatin regulators (*17–19*). At the molecular level, MGO-mediated histone glycation disrupts nucleosome stability, histone-DNA interactions and functional epigenetic marks. More broadly, MGO can modify chromatin- and DNA-associated nuclear proteins, including through DNA-protein crosslinks (*10, 17, 20, 21*). Beyond chromatin, lamin A/C has also been proposed as a target for carbonyl modifications, however direct evidence remains limited, and the mechanical consequences of lamina-associated glycation remain unknown (*22*). Thus, while nuclear glycation is emerging as a key biochemical feature of metabolic stress, it remains unclear how these modifications collectively alter organelle viscoelasticity or dictate how mechanical cues propagate through the nucleoplasm. Crucially, whether such mechanochemical changes regulate downstream mechanosensitive signaling in cells or also manifest in native tissues exposed to metabolic stress remain unaddressed.

Bridging the gap between biochemical modification detection and organelle mechanics requires evaluating how glycated nuclei physically behave under force. Here, we combine atomic force microscopy (AFM) nanoindentation, correlative confocal imaging, 3D intranuclear deformation tracking, targeted nuclear perturbations, and mouse models of metabolic disease to determine how carbonyl stress directly remodels nuclear mechanics and force-sensing across scales. We show that MGO glycation acts directly on load-bearing nuclear structures, shifting the nucleus towards a stiffer, less fluid, and more mechanically constrained state. In live cells, this nucleus-intrinsic material shift persists independently of cytoskeletal tension and biases YAP mechanosensitive signaling. Furthermore, in native cardiac tissue, nuclear stiffening occurs exclusively when metabolic dysfunction is paired with nuclear MGO adduct accumulation. Together, our study establishes non-enzymatic glycation as a direct physical mechanism converting metabolic dysfunction into altered nuclear mechanics and cellular force sensing.

## RESULTS

### MGO glycation directly remodels nuclear mechanics

We first tested whether MGO-derived glycation is sufficient to alter the mechanical properties of the nucleus. To isolate intrinsic nuclear effects from cytoplasmic or cytoskeletal contributions (*23*), we used nuclei isolated from HeLa Kyoto cells stably expressing lamin A/C-Enhanced Green Fluorescent Protein (EGFP) and histone 2B (H2B)-mCherry (**Figure S1a**). This approach enables simultaneous AFM nanoindentation and live confocal visualization of nuclear lamina and chromatin, the major load-bearing nuclear components. Isolated nuclei were incubated *ex-cellulo* with 50 mM MGO for 6 h, while paired controls were measured in phosphate-buffered saline (PBS) alone for the same duration. We used this accelerated treatment at high MGO concentration on purified nuclei as a sufficiency assay to generate detectable MGO-derived adducts (e.g. hydroimidazolone-1 (MG-H1), N^ε^-(carboxyethyl)lysine (CEL), imidazolium crosslink (MODIC)) (*8, 11, 24, 25*) and examine their direct effects on nuclear mechanics, independently of cellular adaptation (**Figure 1a**).

**Fig. 1.**
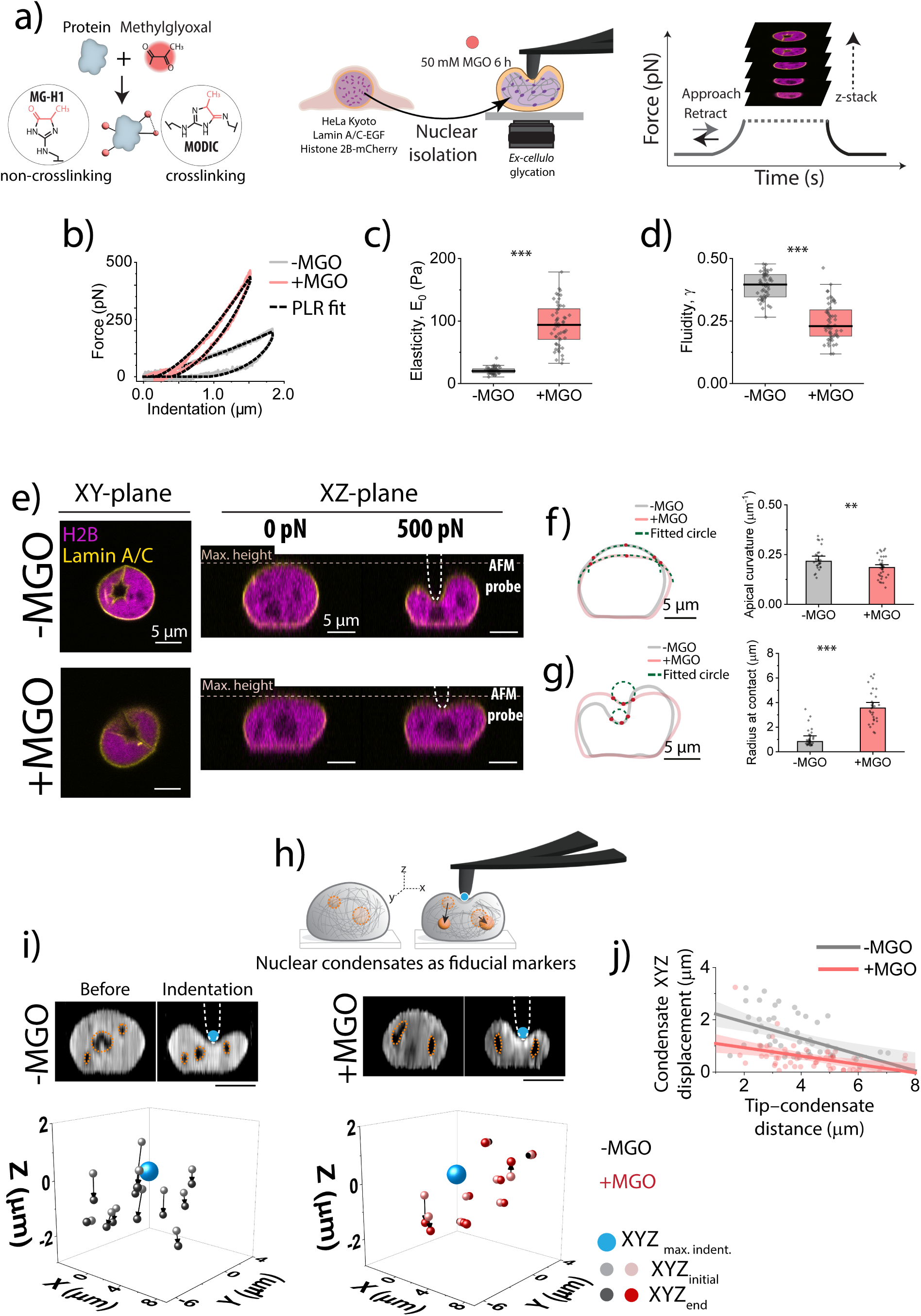
MGO-derived glycation stiffens isolated nuclei, reduces fluidity and restricts intranuclear deformation propagation. **(a)** Experimental setup. Left: MGO reacts non-enzymatically with nucleophilic residues in nuclear proteins, including lysine- and arginine-rich substrates, producing diverse MGO-derived adducts. MG-H1 is shown as an example of a non-crosslinking arginine-derived adduct, whereas MODIC-type chemistry illustrates an MGO-derived lysine-arginine crosslink. Center: isolated HeLa Kyoto lamin A/C–EGFP/H2B–mCherry nuclei were incubated *ex-cellulo* with 50 mM MGO or PBS for 6 h. Right: correlative AFM–confocal measurements were used to quantify nuclear viscoelasticity and visualize force-induced nuclear deformation in 3D. The AFM cantilever approaches the nucleus until a predefined setpoint force is reached, followed by a 60 s hold segment at constant force during which a confocal z-stack is acquired; the cantilever then retracts back to its original position. **(b)** Representative force– indentation curves for −MGO (gray) and +MGO (red) nuclei, with power-law rheology (PLR) fits shown as dashed black lines. **(c)** Quantification of the scaling modulus *E*_0_ and **(d)** fluidity coefficient *γ* for −MGO and +MGO nuclei. Each dot represents the mean of three consecutive indentations per nucleus; boxes indicate 25th–75th percentiles, whiskers indicate outlier boundary and the median of the distribution is represented with a thicker horizontal line. Data were pooled from three independent paired experiments (N(−MGO)=51 nuclei, N(+MGO)=54 nuclei). **(e)** Representative confocal z-stacks of −MGO and +MGO nuclei during AFM indentation. Left: XY plane at the indentation site; right: XZ projections before and after indentation. Lamin A/C is shown in yellow and Histone 2B in magenta. Initial nuclear height (max. height) and the position of the AFM probe are indicated. **(f)** Apical curvature analysis before indentation of nuclei shown in (e), with nuclear contour shown in gray (-MGO) and red (+MGO), fitting limits marked as red dots, and result fitted circle in green. Each point represents the mean curvature from XZ and YZ projections of one nucleus; bars indicate median ± SE. Data are representative of at least three independent experiments (N(−MGO)=29 nuclei, N(+MGO)=35 nuclei). **(g)** Radius at contact analysis upon indentation of nuclei shown in **(e),** with nuclear contour shown in gray (-MGO) and red (+MGO), fitting limits marked as red dots, and result fitted circle in green. Each point represents the mean radius obtained from XZ and YZ projection of one nucleus. Bars show median ± SEM and data were pooled from at least three independent experiments (N(−MGO)=31 nuclei, N(+MGO)=30 nuclei). **(h)** Schematic representation of nuclear condensates used as fiducial markers to quantify intranuclear force propagation during AFM indentation. The 3D distance between the initial condensate position before force application, (open orange circles) and the final position after force application (filled orange circles) was measured for each condensate. The tip– condensate distance was calculated using the AFM tip position at the indentation site (blue). **(i)** Representative condensate displacement in −MGO and +MGO nuclei before and after indentation. Top: confocal z-projections showing condensate position in orange and the AFM tip position in blue; scale bar: 5 µm. Bottom: 3D quantification of condensate positions before force application in lighter shade and after force application in darker shade, with arrows indicating total displacement and the tip position shown as a blue sphere. **(j)** XYZ nuclear condensate displacement as a function of tip–condensate distance during indentation for −MGO and +MGO nuclei. Each point represents a single condensate across two independent experiments (N(-MGO)=50 condensates, N(+MGO)=50 condensates). Linear fits indicate overall trends for -MGO nuclei in gray and +MGO nuclei in red, along with 95% confidence intervals. Statistical analysis was performed using the Mann–Whitney test; *p < 0.05, **p < 0.01, ***p < 0.001; n.s., not significant.

We quantified the mechanical response of individual nuclei by AFM quasi-static force-indentation measurements. Compared with paired controls, MGO-treated nuclei display steeper force-indentation curves and reduced hysteresis, indicating increased resistance to indentation and reduced energy dissipation during the loading cycle (**Figure 1b**). Given the viscoelastic nature of cell nuclei, we analyzed force-indentation curves using a power-law rheology model (*26, 27*). In this framework, we quantified the scaling modulus (*E*₀), representing the apparent stiffness scale, and the power-law exponent (*γ*), which defines the fluidity coefficient of the viscoelastic response. Here, *γ* values closer to 0 indicate a more solid-like response, whereas values closer to 1 reflect increasingly fluid-like behavior. MGO treatment increases the scaling modulus, from *E_0_* = 20.2 ± 0.7 Pa to *E*_0_ = 94 ± 5 Pa (median ± standard error of the mean (SEM), **Figure 1c**) and reduces fluidity from *γ =* 0.396 ± 0.007 to *γ* = 0.23 ± 0.01 (median ± SEM, **Figure 1d**). Time-dependent measurements further show the mechanical response develops progressively during MGO incubation, with the decrease in fluidity preceding the marked stiffness increase (**Figure S1b,c**). Conversely, control nuclei remain mechanically stable throughout the experiment. To further investigate the kinetics of MGO-derived glycation, we reduced MGO to a concentration of 10 mM for 6h. We observe a weaker but significant response in terms of scaling modulus and no changes in fluidity (**Figure S1d,e**), confirming that the observed mechanical remodeling scales with MGO concentration.

To test whether the observed MGO-induced mechanical phenotype could be also caused by buffer-induced osmotic or ionic stress, we exposed isolated nuclei to hyperosmotic high-phosphate PBS (see **Methods, Figure S1f,g**). Consistent with previous reports that osmotic and ionic perturbations can rapidly alter chromatin organization, nuclear size and mechanics (*28, 29*), this condition caused an immediate reduction in fluidity, *γ*, that remained stable over time. However, it did not reproduce the progressive MGO-induced increase in *E*_0_. Thus, osmotic or ionic stress can acutely affect the dissipative response of isolated nuclei, but it does not recapitulate the time-dependent viscoelastic remodeling induced by MGO-driven glycation.

To directly visualize how AFM-applied mechanical forces are translated into nuclear deformation, we applied constant indentation forces while acquiring confocal z-stacks (see **Methods**, **Figure 1e**). Representative confocal images of lamin A/C and chromatin reveal clear differences in nuclear morphology before indentation, with control nuclei exhibiting a more pronounced dome-like apical profile compared to flatter MGO-treated nuclei. We quantified nuclear geometry before force application. MGO-treated nuclei display a less curved apical profile than controls, together with changes in global nuclear dimensions, including increased width and a modest decrease in height (**Figure 1f** and **Figure S2a,b**). AFM indentation produces a sharp local deformation with pronounced height loss in control nuclei, while MGO-treated nuclei require approximately two-fold higher forces to reach comparable indentation depths and show reduced height loss during the hold segment (**Figure 1e**, **Figure S2c–e**). The deformation profile at the AFM contact region is broader and flatter in MGO-treated nuclei, as quantified by an increased radius at contact (**Figure 1g**). The altered apical curvature and broader contact radius observed after MGO treatment point to a change in the mechanical behavior of the nuclear envelope–lamina region. Since lamin A/C is a major load-bearing component of this region, we tested whether it contributes to this geometric response. In lamin A/C knock out (lamin A/C*^KO^*) nuclei, the MGO-induced change in nuclear shape and contact geometry is largely attenuated. Apical curvature remains unchanged and the increase in radius at contact is modest (**Figure S2f-h**).

Having established that MGO-treated nuclei resist local deformation, we further investigated whether this remodeling also changes how deformation propagates through the nuclear interior. This spatial aspect is important because the way the nucleus filters and transmits mechanical inputs can influence chromatin organization, nuclear architecture and mechanosensitive pathways that shape gene regulation (*30, 31*).

We used visible nuclear condensates as internal fiducial markers and tracked their three-dimensional displacement during force application (**Figure 1h** and **Figure S2i,j**). We first quantified the fraction of condensates that underwent detectable displacement at 100 pN and 500 pN. In control nuclei, higher applied force increased the proportion of displaced condensates. In contrast, this force-dependent increase is not observed in MGO-treated nuclei, indicating that glycation reduces propagation of the applied deformation through the nuclear interior (**Figure S2i,j**). Next, we measured condensate displacement as a function of distance from the AFM contact site at comparable indentation depths (see **Methods**). Previous work showed that nuclear deformation can be transmitted to subnuclear structures and that nucleolar displacement can report mechanical propagation into the nucleus (*32–34*). Our results show that control nuclei condensates were displaced by up to ∼3 µm during force application. In contrast, MGO-treated nuclei display an approximately threefold reduction in condensate displacement compared with controls (**Figure 1i,j**).

Together, these results show that MGO glycation directly stiffens the nucleus, reduces its fluidity and limits the propagation of local deformation through the nuclear interior. We next investigated the molecular basis of this mechanical remodeling.

### Molecular origin of the mechanical phenotype

Nuclear mechanics emerge from the coupled contributions of the lamina and chromatin, but whether MGO-derived glycation remodels these load-bearing compartments in a way that explains the mechanical phenotype remains unclear. We therefore mapped the spatial distribution of MGO-derived adducts in purified nuclei and tested how perturbing lamin A/C and chromatin state affects glycation susceptibility and nuclear mechanics (**Figure 2a**). We used two complementary perturbations: lamin A/C*^KO^* to assess the contribution of the lamina, and suberoylanilide hydroxamic acid (SAHA) treatment to increase histone acetylation and reduce the availability of lysine ε-amino groups for reaction with MGO. Lamin A/C depletion and SAHA-induced histone 3 lysine 9 acetylation (H3K9ac) accumulation were confirmed by quantitative immunofluorescence (**Figure 2b,c** and **Figure S3a-c**).

**Fig. 2.**
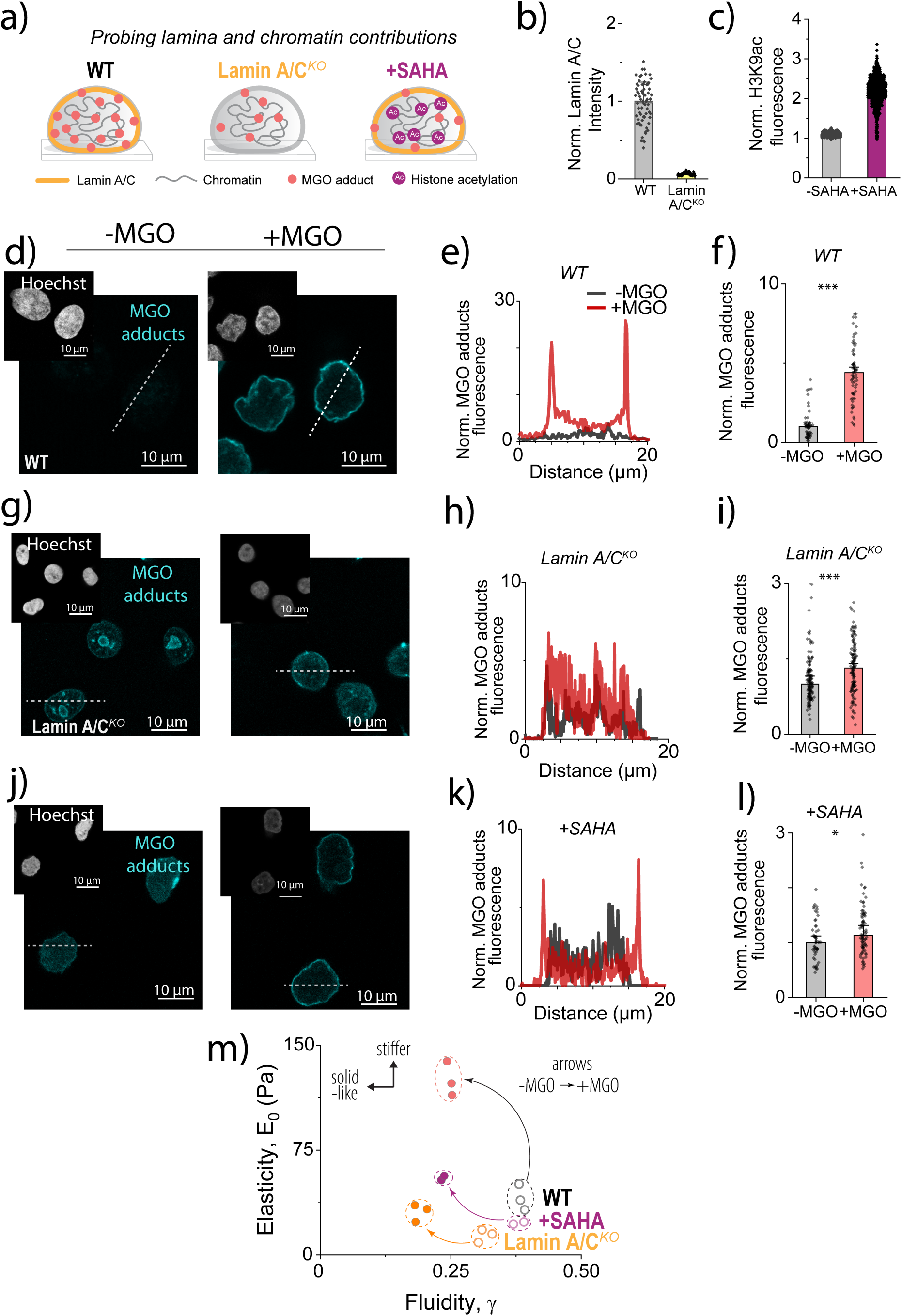
Lamina and chromatin shape MGO-induced nuclear stiffening. **(a)** Schematic illustrating the experimental strategies used to probe how chromatin state and lamin A/C contribute to the mechanical response to MGO-derived glycation. HeLa WT nuclei were compared with SAHA-treated nuclei, used to increase histone acetylation and alter chromatin organization, and with lamin A/C^KO^ nuclei, used to assess the contribution of the nuclear lamina. MGO-derived adducts formed upon incubation with 50 mM MGO for 6 h are indicated in red. **(b)** Quantification of normalized lamin A/C fluorescence in HeLa WT versus lamin A/C^KO^ nuclei, confirming the depletion of lamin A/C protein in knock-out nuclei. Each dot represents one nucleus; bars indicate median ± SEM (N(WT)=67 nuclei, N(lamin A/C^KO^)=82 nuclei) **(c)** Quantification of normalized H3K9ac fluorescence in untreated and SAHA-treated cells, confirming increased histone acetylation after SAHA treatment. Each dot represents one nucleus; bars indicate median ± SEM (N(-SAHA)=1400 cells, N(+SAHA)=885 cells). **(d)** Representative confocal images of control and MGO-treated nuclei stained for MGO-derived protein adducts in cyan. Insets show Hoechst labeling in white. **(e)** Representative line-scan intensity profiles of normalized MGO-adduct fluorescence in −MGO and +MGO nuclei, corresponding to the dashed lines shown in **(d)**. **(f)** Quantification of normalized nuclear MGO-adduct fluorescence. Each dot represents the mean intensity of one nucleus, background-subtracted and normalized to the median of −MGO controls; bars show median ± SEM. Data were pooled from two independent paired experiments (N(-MGO)=62 nuclei, N(+MGO)=68 nuclei). **(g)** Representative confocal images of untreated and MGO-treated lamin A/C^KO^ nuclei stained for MGO-derived protein adducts. Hoechst staining is shown in white in the insets, and MGO adducts are shown in cyan. **(h)** Representative line-scan intensity profiles of normalized MGO-adduct fluorescence in -MGO and +MGO lamin A/C^KO^ nuclei corresponding to the dashed lined in **(g)**. **(i)** Quantification of normalized MGO-adduct fluorescence in lamin A/C^KO^ nuclei. Data were pooled from two independent paired experiments. Each dot represents one nucleus; bars indicate median ± SEM. (N(-MGO)=157 nuclei, N(+MGO)=148 nuclei). **(j)** Representative confocal images of untreated and SAHA-treated nuclei stained for MGO-derived protein adducts. Hoechst staining is shown in white in the insets, and MGO adducts are shown in cyan. **(k)** Representative line-scan intensity profiles of normalized MGO-adduct fluorescence in -MGO and +MGO SAHA-treated nuclei corresponding to the dashed lined in **(j)**. **(l)** Quantification of normalized MGO-adducts fluorescence in SAHA-treated nuclei. Data were pooled from three independent paired experiments. Each dot represents one nucleus: bars indicate median ± SEM; (N(-MGO)=53 nuclei, N(+MGO)=79 nuclei) **(m)** Scaling modulus *E*₀ plotted as a function of the fluidity coefficient *γ* for isolated nuclei from HeLa WT, SAHA-treated WT and lamin A/C^KO^ cells, measured under −MGO and +MGO conditions. Arrows indicate the mechanical shift induced by MGO treatment within each condition. Panels **(d),(g),(j)**: Display ranges were adjusted separately between WT, lamin A/C^KO^ and SAHA datasets due to differences in signal dynamic range; quantitative comparisons are based on measured fluorescence intensities. Each dot represents the median value from an independent biological replicate. Statistical analysis was performed using the Mann–Whitney test; *p < 0.05, **p < 0.01, ***p < 0.001; n.s., not significant.

In wild-type (WT) nuclei, quantitative immunofluorescence shows that MGO treatment increases both the pan-adduct signal (anti-MGO) and MG-H1, a major arginine-derived MGO adduct, by approximately fivefold relative to controls (**Figure 2d-f** and **Figure S4a**). However, these signals show distinct spatial distributions: the pan-adduct signal is enriched at the nuclear periphery, whereas MG-H1 is broadly distributed throughout the nucleus. This suggests that although MGO-derived glycation occurs throughout the nucleus, a subset of MGO-reactive epitopes is enriched or preferentially detected near the nuclear envelope. Omission of the anti-MGO primary antibody abolished the signal, confirming staining specificity (**Figure S4b**). In parallel, lamin A/C immunostaining is markedly reduced in MGO-treated WT nuclei following anti-MGO labelling, compatible with modification or masking of lamin A/C-associated epitopes (**Figure S4c**).

Given its pronounced peripheral enrichment, we used the pan-adduct signal as a spatially resolved readout of nuclear glycation to test whether lamin A/C loss or histone hyperacetylation alters MGO-adduct accumulation and distribution. Untreated nuclei display a low but detectable pan-adduct signal, consistent with basal MGO-derived glycation arising from endogenous metabolism (*10*). In lamin A/C*^KO^* nuclei, MGO still increases the pan-adduct signal, but to a lesser extent compared to WT nuclei, and the sharp peripheral enrichment is largely lost (**Figure 2g-i**). SAHA pre-treatment similarly attenuates MGO-adduct accumulation, although the residual signal remains enriched at the nuclear periphery (**Figure 2j-l**).

We then probed whether these perturbations change nuclear mechanics (**Figure 2m**). Under basal conditions, lamin A/C*^KO^* nuclei are softer (*E*_0_ =13 ± 1 Pa, median ± SEM) than WT nuclei (*E*_0_ =36 ± 2 Pa, **Figure S5a-d**), in line with the load-bearing role of lamin A/C, and also less fluid (*γ* =0.311 ± 0.006 versus *γ* =0.390 ± 0.007). Upon MGO treatment, lamin A/C*^KO^*nuclei stiffen modestly, reaching 32 ± 1 Pa in contrast with WT nuclei, which reach 122 ± 8 Pa. Fluidity decreases to *γ* =0.19 ± 0.006, which is even lower than MGO-treated WT nuclei, which drop to *γ* =0.248 ± 0.009.

SAHA-treated nuclei show a similar separation between stiffness and fluidity responses. Histone hyperacetylation lowers basal nuclear scaling modulus (*E*_0_ = 23 ± 2 Pa, median ± SEM versus *E*_0_ = 36 ± 2 Pa in untreated WT counterparts), without altering fluidity (*γ* = 0.39 ±0.01 versus *γ* =0.390 ± 0.007; **Figure S5a,b,e,f**). Following MGO treatment, SAHA-treated nuclei reach only 54 ± 3 Pa, substantially less than MGO-treated WT nuclei (*E*_0_ = 122 ± 8 Pa), indicating that histone hyperacetylation attenuates the stiffening response. In contrast, MGO reduces fluidity to a similar extent in SAHA-treated and WT nuclei (*γ* = 0.23 ± 0.01, compared to *γ* =0.248 ± 0.009 for WT nuclei; **Figure S5e,f**). Importantly, SAHA-treated HeLa cells and isolated nuclei show the same mechanical trend relative to their respective controls, with reduced scaling modulus and unchanged fluidity (**Figure S5g,h**). This agrees with previous work showing that chromatin decompaction induced by histone acetylation softens the nucleus (*35*).

These perturbations show that glycation slows the stress-relaxation response even when lamina- or chromatin-dependent stiffening is attenuated. We next asked whether chronic carbonyl stress produces a related nuclear mechanical phenotype in living cells, where the nucleus remains coupled to the cytoskeleton and cellular adaptation pathways could either amplify or buffer the effect of glycation.

### Chronic carbonyl stress induces nucleus-intrinsic stiffening

Intracellular MGO levels remain broadly difficult to define due to its high reactivity and challenging quantification (*36*), however estimates suggest that in type 2 diabetes, the intracellular concentration could sit in the micromolar range (*37*). We first sought to define a threshold for chronic MGO exposure compatible with HeLa cell survival. We therefore performed cell cytotoxicity and reactive oxygen species (ROS) assays at concentrations ranging between 100 µM – 500 µM for 72h (Methods, **Figure S6**). At 500 µM, MGO causes only a moderate reduction in mitochondrial activity and does not markedly increase ROS levels (**Figure S6a,b**). We therefore used 500 µM MGO for 72 h in subsequent cell-based experiments (**Figure 3a**). Confocal microscopy reveals irregular nuclear contours, local lamin A/C–positive accumulations, discontinuities at the nuclear periphery, and occasional nuclear blebs in MGO-treated cells (**Figure 3b**).

**Fig. 3.**
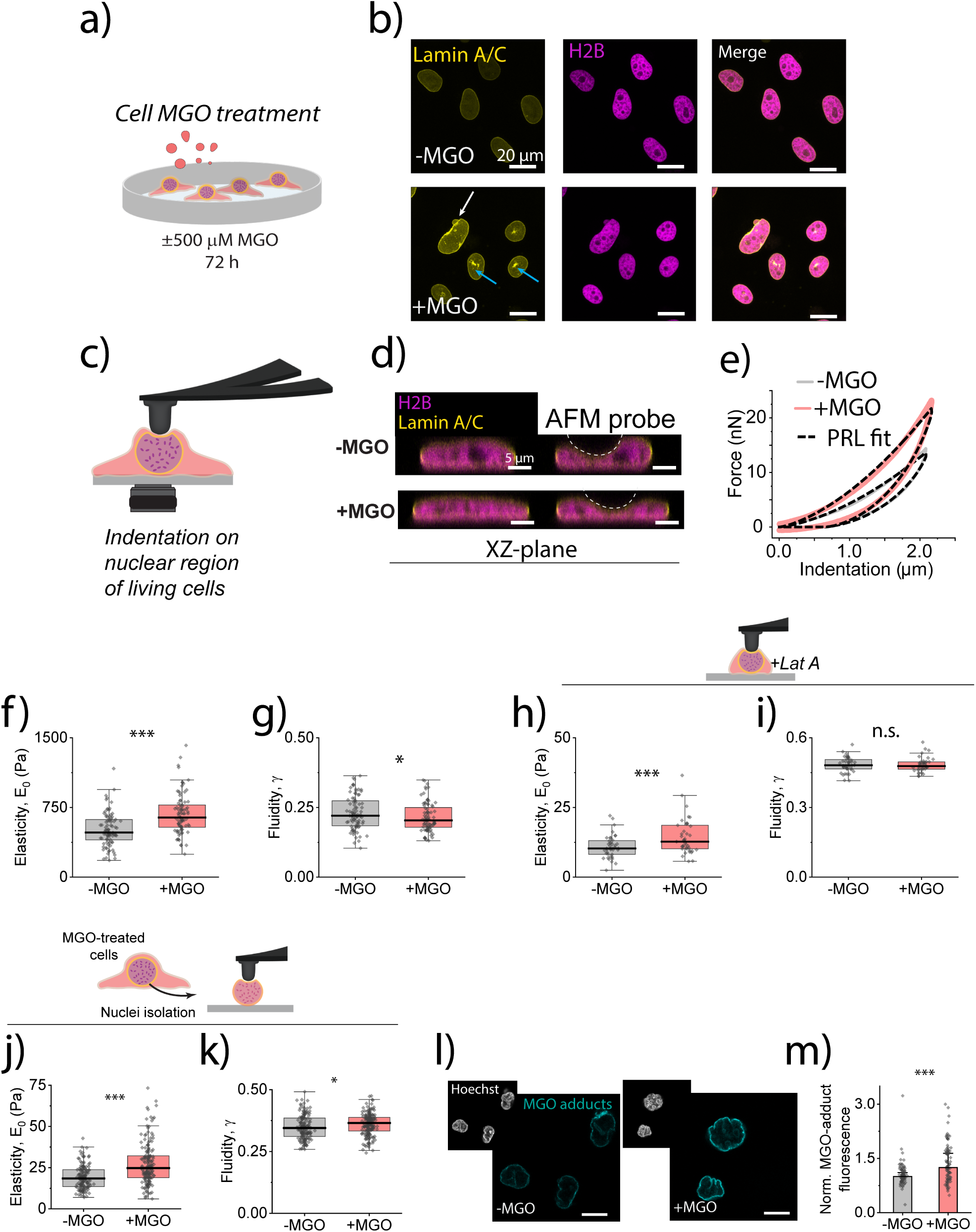
Chronic MGO exposure alters nuclear-envelope organization and induces nucleus-intrinsic stiffening *in-cellulo*. **(a)** Experimental workflow. HeLa Kyoto cells stably expressing lamin A/C–EGFP and Histone H2B–mCherry were cultured in complete medium supplemented or not with 500 µM MGO for 72 h, followed by live-cell AFM–confocal analysis and post-treatment nuclear isolation. **(b)** Representative confocal images of control (−MGO) and MGO-treated (+MGO) living cells. Lamin A/C is shown in yellow and Histone H2B in magenta. Chronic MGO exposure is associated with altered nuclear morphology (white arrows) and discontinuous or irregular lamin A/C signal (blue arrows), consistent with altered nuclear-envelope organization. **(c)** Schematic of AFM indentation targeted to the nuclear region of living cells. Force–indentation curves were acquired on top of the nucleus using a spherical AFM probe with a 5 µm radius. **(d)** Representative confocal XZ projections of −MGO and +MGO cells before and during AFM indentation. The dashed white line indicates the shape of the indenting AFM probe. **(e)** Representative force–indentation curves acquired over the nuclear region of −MGO and +MGO cells. Dashed black lines indicate PLR fits. **(f)** Quantification of the scaling modulus *E*_0_ and **(g)** fluidity coefficient *γ* measured over the nuclear region. Each dot represents the mean of three consecutive indentations per cell; boxes indicate 25th–75th percentiles, whiskers indicate outlier boundary and the median of the distribution is represented with a thicker horizontal line. Data were pooled from four independent biological replicates (N (-MGO)= 87 cells, N(+MGO)= 84 cells). **(h)** Quantification of the scaling modulus *E*_0_ and **(i)** fluidity coefficient *γ* measured over the nuclear region after treatment with 1 µM latrunculin A for 20 min. Nuclear-region stiffening persists after actin disruption, whereas *γ* shows no significant change. Each dot represents the mean of three consecutive indentations per cell; boxes indicate 25th–75th percentiles, whiskers indicate outlier boundary and the median of the distribution is represented with a thicker horizontal line. Data were pooled from two independent biological replicates (N(-MGO)=40 cells, N(+MGO)=39 cells). **(j)** Quantification of the scaling modulus *E*_0_ and **(k)** fluidity coefficient *γ* of nuclei isolated from −MGO and +MGO cells. Chronic cellular MGO exposure causes persistent nuclear stiffening after isolation, whereas changes in fluidity remain modest. Each dot represents the mean of three consecutive indentations per nucleus; boxes indicate 25th–75th percentiles, whiskers indicate outlier boundary and the median of the distribution is represented with a thicker horizontal line. Data were pooled from six independent biological replicates (N(-MGO)=145 nuclei, N(+MGO)=167 nuclei). **(l)** Representative confocal images of nuclei stained for MGO-modified proteins. MGO adducts are shown in cyan; Hoechst is shown in white in the insets. **(m)** Quantification of nuclear MGO-adduct fluorescence in isolated nuclei from −MGO and +MGO cells. Each dot represents the mean intensity of one nucleus after background subtraction and normalization to the median of −MGO controls; bars indicate median ± SEM. Data were pooled from three independent biological replicates (N(-MGO)=86 nuclei, N(+MGO)=91 nuclei). Statistical significance was evaluated using two-sided Mann–Whitney tests; *P < 0.05, **P < 0.01, ***P < 0.001; n.s., not significant.

We next evaluated if these changes in nuclear-envelope organization are accompanied by altered nuclear-region viscoelastic properties. Probing the nuclear region of living cells by AFM while monitoring nuclear deformation by confocal microscopy allowed us to confirm the indentation range needed to deform the underlying nucleus (around 2 µm, **Figure 3c,d**). Power-law rheology analysis of the recorded force-indentation curves shows that MGO increases the scaling modulus *E*_0_ over the nuclear region from 510 ± 19 Pa to 680 ± 24 Pa (median ± SEM), while changes in the fluidity coefficient *γ* are modest compared to the strong loss observed upon direct *ex-cellulo* glycation (*in-cellulo*: 0.221 ± 0.006 to 0.203 ± 0.006 after MGO exposure, **Figure 3e-g**).

Since the actin cytoskeleton contributes to nuclear-region mechanics, transmits prestress and may be a direct target of MGO glycation (*38, 39*), we examined whether MGO-induced stiffening persists after cytoskeletal disruption. Cells were pre-incubated with 500 µM MGO for 72 h, followed by treatment with 1 µM latrunculin A (Lat A) for 20 minutes prior to AFM measurements. Lat A strongly softens and fluidizes the nuclear region of both control and MGO-treated, reflecting the loss of actin-dependent prestress and mechanical coupling to the nucleus. However, MGO-treated cells remain 30% stiffer than matched controls even after actin depolymerization, a relative increase mirroring that observed in Lat A-untreated cells (**Figure 3h,i**). Fluidity is not significantly altered under these conditions. We conclude that chronic MGO exposure causes intrinsic nuclear-region stiffening that persists independently of the actin cytoskeleton.

To confirm that this observed phenotype stems from intrinsic nuclear glycation, we performed AFM nanoindentation on nuclei isolated from control and MGO-treated cells. The MGO-treated nuclei are significantly stiffer, with *E*_0_ increasing from 18.4 ± 0.8 Pa to 25 ± 1 Pa (median ± SEM), while *γ* exhibits only modest changes, in line with our living cells observations (**Figure 3j,k**). Furthermore, these nuclei show a marked accumulation of MGO adducts compared with paired controls (**Figure 3l,m**).

Collectively, these findings suggest that chronic glycation stress directly targets the nucleus of living cells, triggering a stable, structural stiffening response that persists outside the cellular environment.

### MGO-induced nuclear remodeling biases YAP mechanosensitivity

Mechanosensitive signaling relies on the physical properties of the nucleus and its structural coupling to the cytoskeleton. To test whether MGO-induced nuclear stiffening alters cellular mechanosensing, we evaluated the nucleocytoplasmic partitioning of Yes-associated protein (YAP), a mechanosensitive transcriptional co-regulator. As a downstream effector of the Hippo pathway, YAP shuttles dynamically between the cytoplasm and the nucleus, enabling cells to translate physical forces into gene-expression programs (*40*). On soft substrates, reduced cell spreading and lower actomyosin tension generally limit YAP nuclear accumulation, whereas stiff substrates promote spreading, nuclear deformation and YAP enrichment in the nucleus (*41*).

To determine if chronic MGO treatment alters this substrate stiffness-dependent response, control and MGO-treated (500 µM, 72 h) HeLa cells were seeded onto soft or stiff polyacrylamide hydrogels, with Young’s moduli of 0.1 kPa and 50 kPa, respectively (**Figure 4a**). On soft hydrogels, cells maintain a rounded morphology, whereas on stiff gels they adopt a highly spread morphology regardless of treatment (**Figure 4b,c**). In control cells, the YAP nuclear-to-cytoplasmic (N/C) ratio increases from soft to stiff substrates, confirming the expected stiffness-dependent YAP translocation (**Figure 4d,f**). In contrast, MGO treatment significantly elevates nuclear YAP accumulation on both substrates (**Figure 4d,f**). We observe the strongest effect on stiff hydrogels, where MGO-treated cells exhibit an approximately 50% increase in the YAP N/C ratio compared to untreated controls. A smaller but statistically significant 10% increase is also detected on soft hydrogels, where actomyosin tension is otherwise low.

**Fig. 4.**
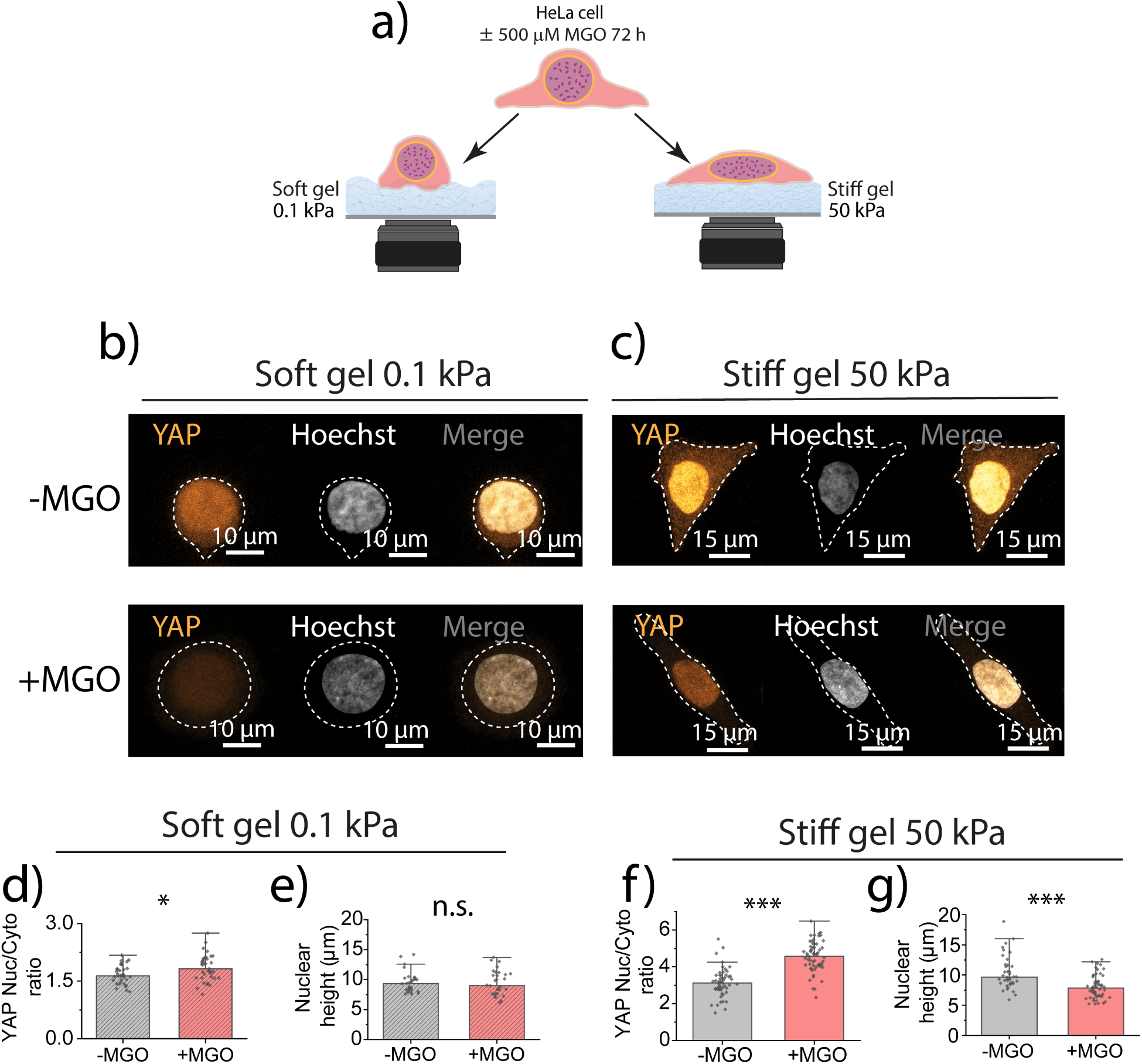
Chronic MGO exposure biases stiffness-dependent YAP nuclear localization. **(a)** Schematic representation of HeLa cells cultured for 72 h in complete medium with or without 500 µM MGO and then plated on soft or stiff polyacrylamide gels to modulate substrate stiffness. Soft gels had a Young’s modulus of 0.1 kPa, whereas stiff gels had a Young’s modulus of 50 kPa. Representative confocal images of control (−MGO) and MGO-treated (+MGO) cells grown on soft **(b)** or stiff **(c)** substrates and stained for YAP and Hoechst. Maximum intensity projections are shown, with YAP in gold and Hoechst in white. Dashed outlines indicate cell boundaries obtained by image thresholding. **(d)** Quantification of the YAP nuclear-to-cytoplasmic ratio in −MGO and +MGO cells on soft 0.1 kPa substrate (N(-MGO)=36 cells, N(+MGO)=37 cells). **(e)** Quantification of nuclear height in -MGO and +MGO cells on soft substrates (N(-MGO)=30 cells, N(+MGO)=30 cells). **(f)** Quantification of the YAP nuclear-to-cytoplasmic ratio in −MGO and +MGO cells on stiff 50 kPa substrate. Each dot represents one cell; bars indicate median ± SEM. Data were pooled from three independent paired biological replicates (N(-MGO)=53 cells, N(+MGO)=60 cells). **(g)** Quantification of nuclear height in -MGO and +MGO cells on stiff substrates (N(-MGO)=39, N(+MGO)=60). Data were pooled from three independent biological replicates. Each dot represents one cell, and bars indicate median ± SEM. Statistical significance was assessed using Mann– Whitney tests; *P < 0.05, ***P < 0.001.

Because YAP localization is sensitive to cell and nuclear geometry, we also evaluated whether the MGO-dependent increase in nuclear YAP could be driven by macroscopic shape changes. We observe that MGO treatment alters cell morphology in a substrate-dependent manner (**Figure 4e,g** and **Figure S7**). On stiff gels, MGO-treated cells display reduced height and apical curvature, indicating increased cell flattening. However, on soft gels, where MGO treatment also significantly elevates the YAP N/C ratio, cell height and apical curvature remain unchanged compared to controls.

Therefore, although MGO promotes cell flattening on stiff substrates, macroscopic geometry alone does not account for the increased nuclear localization of YAP across both mechanical environments. Chronic MGO exposure shifts cellular YAP localization toward the nucleus on both stiff and soft substrates, with the strongest effect on stiff matrices, linking carbonyl stress-induced nuclear remodeling to altered mechanosensitive regulation.

### Nuclear MGO-adduct accumulation predicts solid-like transition across models

To test whether carbonyl stress remodels nuclear mechanics in disease-relevant tissues, we turned to cardiac nuclei from mouse models of metabolic dysfunction. This setting is particularly relevant because diabetes and metabolic syndrome are associated with myocardial stiffening and diastolic dysfunction, and work from our group and others have established that carbonyl stress contributes to mechanical remodeling of cardiac contractile proteins (*6, 8, 38*). We compared two distinct mouse models of metabolic stress: first, a Western Diet (WD) model, in which mice exposed to a high-fat, high-sugar diet develop hyperglycemia and insulin resistance that recapitulate human type 2 diabetes (*42–44*); and second, leptin-deficient (*ob*/*ob)* mice, a canonical genetic model of obesity and severe metabolic dysfunction (*45*). Cardiac nuclei were isolated from each model and matched controls, then analyzed by AFM nanoindentation and confocal imaging (**Figure 5a**). AFM nanomechanical measurements reveal a model-dependent divergence in nuclear mechanical properties. Cardiac nuclei from WD hearts are significantly stiffer than those from chow-fed controls, with basal *E*_0_ increasing by approximately 60% (from 88 ± 9 Pa to 137 ± 9 Pa, median ± SEM; **Figure 5b,d** and **Figure S8a,b**). In contrast, cardiac nuclei from *ob*/*ob* mice exhibit no significant change in *E*_0_ compared with their lean controls (**Figure 5c,d** and **Figure S8g,h).** Nuclear fluidity (*γ*) remains largely unchanged across both models (**Figure 5e** and **Figure S8c,d,i,j**).

**Fig. 5.**
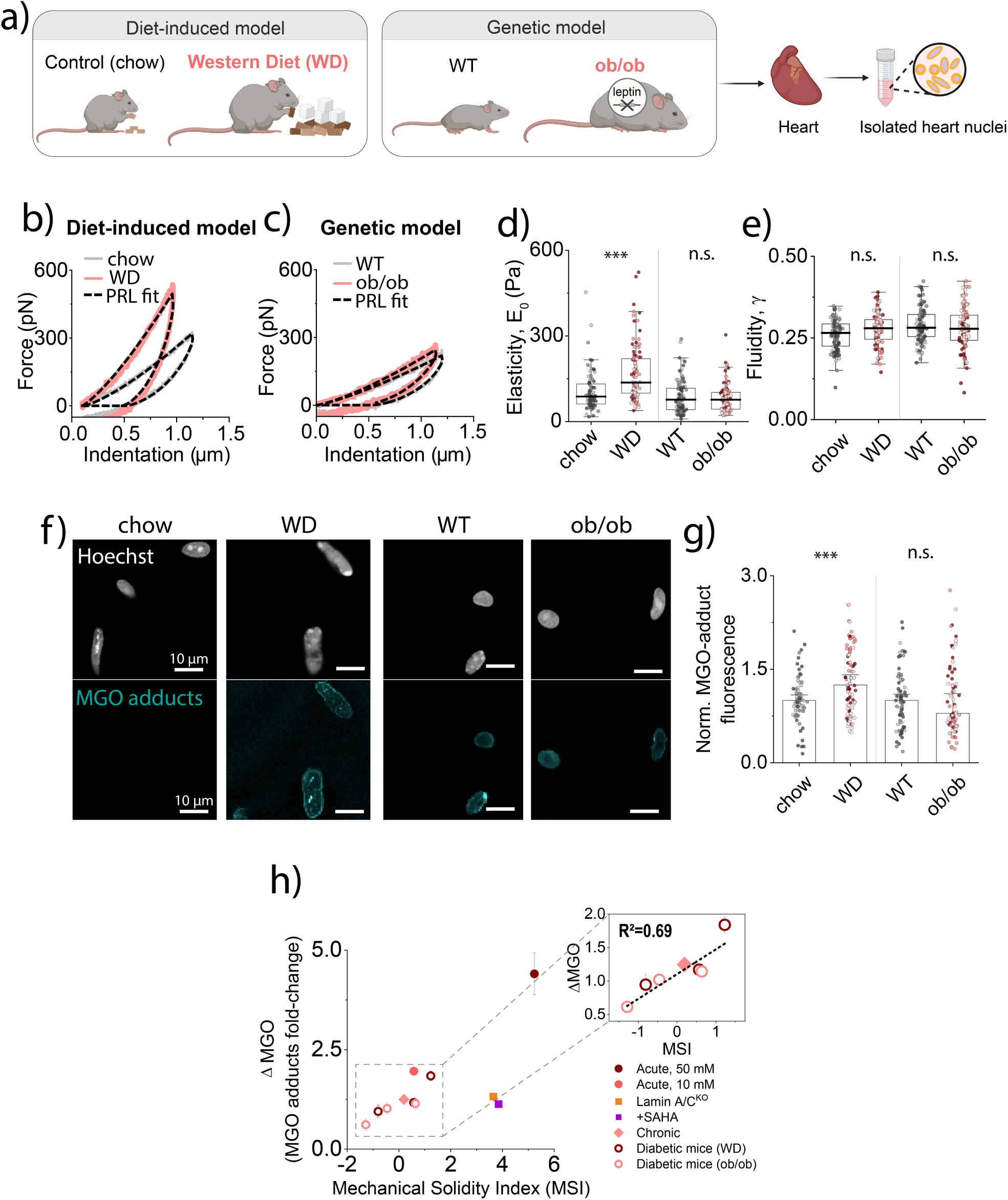
Cardiac nuclear stiffening in metabolic disease tracks with nuclear MGO-adduct accumulation. **(a)** Cardiac nuclei were isolated from two mouse models of metabolic dysfunction and their respective controls: a diet-induced model comparing Western diet (WD) with chow-fed mice (control), and a genetic model comparing leptin-deficient *ob/ob* mice with WT mice. Isolated heart nuclei were then analyzed by AFM force spectroscopy and confocal imaging. **(b)** Representative force–indentation curves from isolated cardiac nuclei obtained from control and WD mice, or **(c)** WT and *ob/ob* mice. Dashed black lines indicate power-law rheology (PLR) fits. **(d)** Quantification of the scaling modulus *E*0 and **(e)** fluidity coefficient *γ* for isolated cardiac nuclei from diet-induced and genetic diabetes models. Each dot represents the mean of three consecutive measurements per nucleus. Nuclei coming from the same animal are represented with a different shade of gray (for control specimens of each model) or in red (for diabetic individuals of each model). Boxes indicate 25th–75th percentiles, whiskers indicate outlier boundary and the median of the distribution is represented with a thicker horizontal line (N(chow)=89 nuclei, N(WD)=104 nuclei, N(WT)=98 nuclei, N(*ob/ob*)= 97 nuclei). **(f)** Representative confocal images of isolated cardiac nuclei from control, WD, WT and *ob/ob* mice stained for MGO-modified proteins and Hoechst. MGO adducts are shown in cyan and Hoechst in white. **(g)** Quantification of nuclear MGO-adduct accumulation in cardiac nuclei from diet-induced and genetic metabolic dysfunction models. Each dot represents the mean normalized fluorescence of a single nucleus (N(chow)=72 nuclei, N(WD)=87 nuclei, N(WT)=73 nuclei, N(*ob/ob*)= 73 nuclei). **(h)** Relationship between nuclear MGO-adduct accumulation and MSI across the experimental models analyzed in this study. MGO-adduct accumulation is expressed as the fold change in nuclear MGO-adduct fluorescence relative to the matched control. MSI was calculated as a signed standardized shift toward increased scaling modulus, *E*_0_, and decreased fluidity, *γ*, relative to each condition’s paired baseline. Higher MSI values indicate a shift toward a stiffer and less fluid, more solid-like nuclear state, whereas values near or below zero indicate little or no solid-like shift. The inset shows the relationship between nuclear MGO-adduct accumulation and MSI for the chronic cellular and cardiac mouse models; the dashed line indicates the linear fit for this subset. Statistical significance was assessed using Mann–Whitney tests; *P < 0.05, ***P < 0.001; n.s., not significant.

As previously, we also quantified nuclear MGO-adduct accumulation across the corresponding preparations (**Figure 5f,g** and **Figure S8e,f,k-n**). WD cardiac nuclei display a significant 25% increase in MGO-adduct signal relative to chow controls (**Figure 5g**), while *ob*/*ob* cardiac nuclei show no detectable increase compared with their matched controls. This direct correspondence between detectable nuclear MGO adducts accumulation and elevated *E*_0_ indicates that nuclear stiffening is not a generic consequence of metabolic dysfunction, but it specifically tracks with immunodetected MGO-adduct burden, consistent with an MGO-driven contribution.

To capture viscoelastic remodeling across systems with different baseline mechanics, we established a *Mechanical solidity index* (MSI). The MSI integrates the standardized changes in scaling modulus *E*_0_ and fluidity coefficient *γ* relative to control variability, where positive values denote solid-like remodeling and negative values reflect fluid-like softening (see **Methods** and **Figure S9**). Across unperturbed carbonyl-stress models, MSI increases with nuclear MGO-adduct burden, linking direct nuclear glycation, chronic cellular exposure and native cardiac tissue within a common solid-like remodeling trajectory (**Figure 5h**). Lamin A/C*^KO^* and SAHA-treated nuclei deviate from this trend. In these conditions, nuclei start from a mechanically softer basal state, and MGO glycation produces a pronounced relative shift toward solid-like behavior, even though their final absolute *E*_0_ remains lower than MGO-treated WT nuclei. This separation is most apparent in fluidity, which decreases sharply despite a reduced MGO-adduct burden and attenuated stiffening. Thus, MSI efficiently captures how far each condition shifts from its own basal mechanical state towards a more solid-like response, linking MGO burden and mechanical remodeling across scales.

## DISCUSSION

Carbonyl stress is a central consequence of diabetes, ageing, and metabolic dysfunction, and has long been linked to the progressive stiffening of cells and tissues (*10, 46, 47*). In the heart, this mechanical remodeling has been explained mainly by extracellular matrix AGEs accumulation and by glycation or oxidative modification of sarcomeric and cytoskeletal proteins (*6, 8, 38, 48*). Yet carbonyl-induced modifications are not restricted to extracellular or cytoplasmic proteins. Several nuclear targets of carbonyl stress have also been identified: MGO-derived histone glycation alters histone–DNA interactions, chromatin accessibility and higher order chromatin architecture, and MGO can also induce DNA–protein crosslinks involving chromatin- and DNA-associated proteins (*17, 19, 21*). Lamin A/C has also been proposed as a putative target of MGO-induced glycation modification in human vascular tissue (*22*). However, whether these modifications alter the physical behavior of the nucleus as a load-bearing material has remained unknown.

In this work, HeLa Kyoto cells expressing lamin A/C–EGFP and H2B–mCherry provided a tractable model for correlative AFM–confocal dissection of the load-bearing nuclear compartments. The concordance between *ex-cellulo* glycated HeLa nuclei and WD cardiac nuclei additionally supports the presence of a shared chemistry-to-mechanics axis. We provide the first evidence that MGO-derived glycation represents a non-enzymatic mechanism by which metabolic stress can remodel nuclear material properties outside the usual enzyme-controlled pathways that regulate lamina and chromatin mechanics (**Figure 6**).

**Fig. 6.**
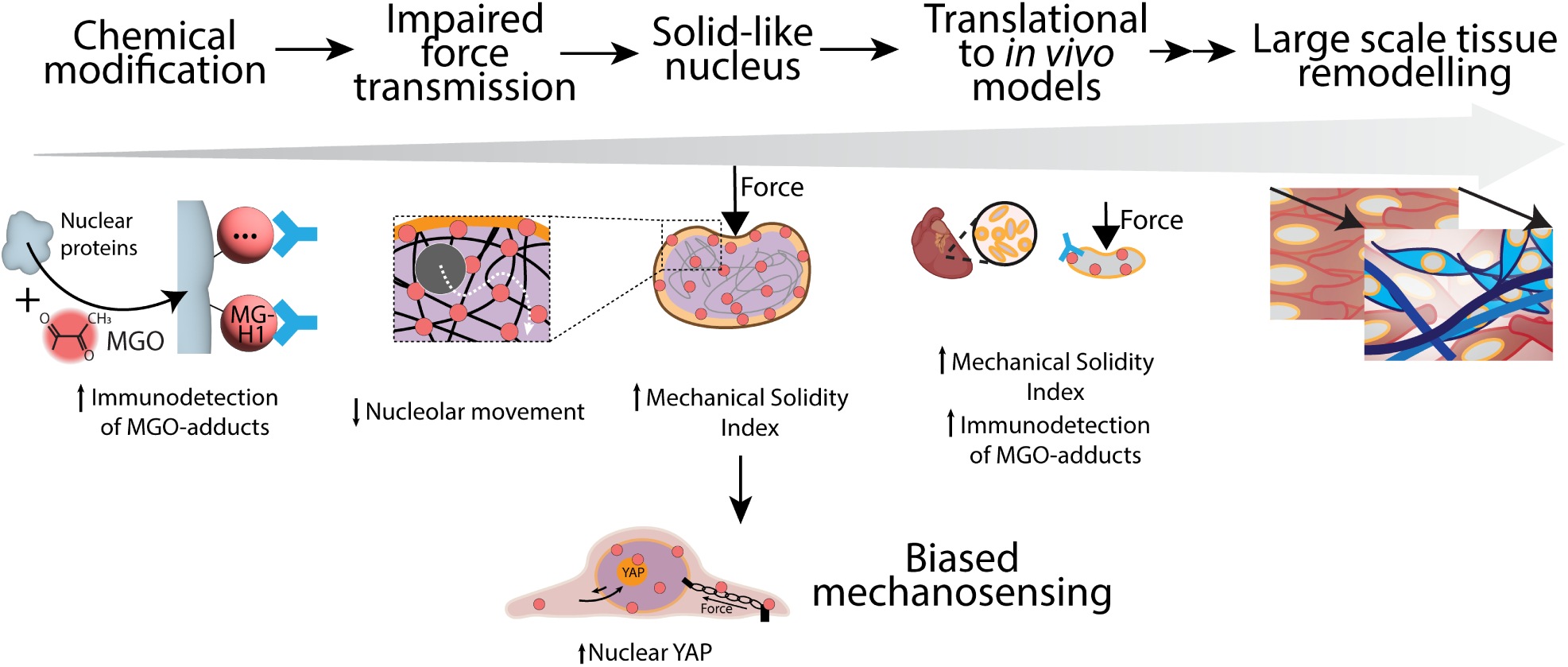
Working model: Carbonyl stress driven by MGO promotes the accumulation of MGO-derived adducts in the nucleus, detected by pan-MGO and MG-H1 immunostaining. This heterogeneous glycation burden is proposed to act on lamina- and chromatin-associated nuclear structures, increasing nuclear stiffness, reducing fluidity and restricting intranuclear deformation propagation. The combined shift toward a more solid-like nuclear state is captured by the MSI. In cells, chronic MGO exposure is associated with increased YAP nuclear localization, while in cardiac nuclei from WD mice, increased MGO-adduct burden is accompanied by elevated MSI, supporting the relevance of this nucleus-intrinsic mechanical remodeling in metabolic disease.

Nuclear mechanics has traditionally been interpreted through lamin mutations or expression changes, chromatin compaction, cytoskeletal prestress, osmotic compression, and enzymatically controlled post-translational modifications such as lamin phosphorylation or histone acetylation (*14*). Non-enzymatic glycation differs from these mechanisms because it is not catalyzed by a dedicated enzyme and is not spatially programmed by the cell. Instead, it depends on local carbonyl burden, exposure time, detoxification capacity and the chemical susceptibility of nuclear proteins. Our findings therefore place non-enzymatic glycation alongside canonical enzyme-driven regulation as a determinant of nuclear material state. This adds a distinct layer to nuclear mechanobiology, as nuclear mechanics may therefore reflect not only rapid and reversible tuning of lamina and chromatin, but also slower covalent modifications that accumulate under carbonyl stress and can leave a persistent mechanical imprint.

The *ex-cellulo* experiments establish that the nucleus itself is mechanically susceptible to MGO-derived glycation. Direct MGO exposure does not simply produce a generic “stiffer nucleus” phenotype. Instead, glycated nuclei resist indentation more strongly, show slower stress relaxation, and deform with a broader, flatter contact geometry characterized by an increased radius at contact and reduced height loss under load. The reduction in internal condensate displacement adds a spatial dimension to this phenotype: following glycation, deformation propagates less efficiently through the nucleoplasm.

Immunofluorescence mapping suggests that the mechanical response to MGO is accompanied by chemically and spatially heterogeneous glycation. MG-H1, a major non-crosslinking arginine-derived MGO adduct, is broadly distributed throughout the nucleus, indicating widespread modification of nuclear substrates. By contrast, the pan-MGO signal is enriched near the nuclear periphery, suggesting that a subset of MGO-derived epitopes accumulates preferentially at, or becomes more detectable near, lamina-associated structures. Non-crosslinking AGE, including MG-H1 and CEL, can alter charge distribution, protein recognition, electrostatic interactions and histone–DNA contacts, while potentially competing with enzymatic PTM sites (*11, 21, 24, 25, 49–52*). Crosslinking or multivalent MGO-derived AGE species, such as MOLD, or MODIC or methylglyoxal derived dihydroxyimidazolidine hemiacetal crosslink (MIDAL) (*53*), although potentially less abundant, could have disproportionate mechanical effects by imposing steric and covalent constraints on the relative motion of lamina-associated proteins, chromatin fibers or neighboring nuclear polymers (*8, 11, 24, 25, 50*). Identifying which specific modifications contribute most to this nuclear mechanical phenotype remains challenging (*13*), as MGO-derived glycation generates a heterogeneous ensemble of AGEs on diverse nuclear targets. Antibody-based detection can map where MGO-derived epitopes accumulate, but it cannot resolve the ratio of non-crosslinking adducts to crosslinking species, the latter being notoriously difficult to quantify by unbiased proteomic approaches. Nevertheless, quantitative LC-MS/MS profiling has mapped extensive MGO-derived adducts across histone tails and nuclear structural networks, establishing nucleosomal and nuclear structural proteins as prime targets for dicarbonyl stress (*17, 54, 55*). The physical crosslinking and charge-neutralization caused by these identified adducts align with our observed viscoelastic shift toward elevated scaling modulus and restricted fluid-like relaxation, providing a molecular basis for the restricted 3D intranuclear deformation observed in our tracking assays. In this way, mechanical probing resolves what chemical profiling alone cannot: it integrates a spatially and chemically complex adduct landscape into a unified, organelle-level material transformation.

Modulating lamin expression and chromatin state pinpoints the structural features that mediate MGO-induced stiffening. Lamin A/C loss reduces peripheral pan-adduct accumulation and attenuates stiffening, while SAHA-induced histone hyperacetylation similarly dampens the stiffening response. These results support the established view that lamina and chromatin are major load-bearing systems of the nucleus and extend this framework to non-enzymatic metabolic chemistry. However, neither perturbation prevents the decrease in the fluidity coefficient, indicating that while glycation-induced stiffening depends strongly on lamina- and chromatin-associated architecture, reduced fluidity likely emerges at higher levels of nuclear glycation, when modifications become sufficiently extensive to constrain molecular rearrangements across the nuclear interior (*21*).

This separation between scaling modulus and fluidity also helps explain why the response differs across *ex-cellulo*, cellular and tissue models. The acute 50 mM MGO treatment is an accelerated chemical test of sufficiency, not a physiological exposure model. Nevertheless, such incubations generate canonical MGO-derived AGEs products, including MG-H1, CEL and MGO-derived crosslinks, that overlap with AGE chemistries detected in biological tissues (*49, 51, 56*). When the MGO concentration is lowered *ex-cellulo*, nuclei still stiffen, but the decrease in fluidity is no longer observed. This is consistent with previous work showing that MGO effects on chromatin architecture depend on the extent of histone glycation: limited modification can alter histone–DNA interactions, whereas stronger or prolonged exposure promotes histone crosslinking and chromatin compaction (*21*). In our system, this suggests that stiffness can arise from a lower or more localized adduct burden, whereas reduced fluidity may require broader modification of chromatin- and lamina-associated substrates, including crosslinking or multivalent constraints that limit molecular rearrangements. In cells, where adducts accumulate slowly during the 72h treatment and in the presence of detoxification pathways, nuclear–cytoskeletal coupling and cellular adaptation, the phenotype is therefore more selective, dominated by nuclear stiffening with modest changes in fluidity. This stiffening is not explained solely by cytoskeletal prestress, as it persists after actin disruption, and nuclei isolated from MGO-treated cells retain both increased stiffness and elevated MGO-adduct accumulation.

The same relationship is evident in native cardiac tissue. Diabetic and metabolically stressed tissues are mechanically remodeled at the extracellular and cytoskeletal levels (*5*), and nuclear mechanics can adapt to these environments through changes in lamin A/C levels, lamin turnover and cytoskeletal coupling. A stiffer diabetic tissue could therefore be expected to produce stiffer nuclei through chronic mechanical adaptation alone. Instead, cardiac nuclear stiffening is detected only in the WD model, where nuclei show increased MGO-reactive adduct signal, and not in *ob/ob* mice, where nuclear MGO-adducts burden does not increase. This lack of stiffening is consistent with the metabolic trajectory of the *ob/ob* model, in which hyperglycemia can be transient or compensated despite persistent obesity and hyperinsulinemia (*57*). Thus, the mechanical phenotype retained after nuclear isolation is not a generic consequence of obesity, diabetic-like metabolism or tissue-level stiffness, but tracks with local nuclear carbonyl-adduct accumulation.

To compare glycation-associated mechanical remodeling across experimental systems with very different basal mechanics, we used the MSI, moving beyond a simple *soft-to-stiff* description based on absolute values. MSI therefore captures how far each model shifts from its own baseline, by combining the increase in *E*_0_ and the decrease in *γ* into a single score. Thus, a large mechanical response in a relatively soft system is not underestimated simply because its final stiffness remains lower than that of a model with a higher basal stiffness. Across unperturbed models, MSI scales with nuclear MGO-adduct burden, supporting a direct relationship between glycation load and the magnitude of solid-like nuclear remodeling. Residual variability likely reflects the fact that immunodetection captures only a subset of the MGO-derived modifications contributing to the mechanical phenotype. Lamin A/C*^KO^* and SAHA-treated nuclei deviate from this relationship, suggesting that both the nuclear envelope and chromatin modulate the mechanical response to glycation.

The relevance of this retained nuclear mechanical phenotype is supported by previous links between carbonyl stress, diabetes and YAP signaling. MGO-induced carbonyl stress has been reported to promote sustained YAP/TAZ nuclear localization and activity in cancer cells through Hsp90 glycation and impaired LATS1 regulation (*47*). In cardiovascular models, diabetes and high glucose have also been associated with YAP activation, including increased nuclear YAP in diabetic cardiac tissue and enhanced YAP activation under high-glucose laminar-flow conditions in endothelial cells (*58, 59*). Our results add a nuclear-mechanical dimension to these observations. Chronic MGO exposure shifts YAP localization toward the nucleus across soft and stiff substrates, with the strongest effect on stiff matrices. Since cell height and apical curvature remain unchanged on soft hydrogels, this response is unlikely to reflect geometry-driven YAP activation alone. Rather, the altered YAP response suggests that MGO-induced nuclear stiffening modifies how matrix stiffness is coupled to YAP nucleocytoplasmic partitioning.

Collectively, our findings establish nuclear glycation as a previously unrecognized process that remodels nuclear viscoelasticity. By altering physical properties that govern force transmission and nuclear compliance, non-enzymatic glycation converts transient metabolic stress into persistent changes in cellular mechanical behavior. This positions the nucleus as an active physical driver of pathology rather than a passive biomarker of metabolic dysfunction. Limiting nuclear carbonyl damage or preserving intrinsic nuclear fluidity may therefore represent a distinct therapeutic strategy to protect organelle function in aging and diabetes.

## MATERIALS AND METHODS

### Methylglyoxal purification

MGO was purchased as a 40% aqueous solution (M0252, Sigma-Aldrich) and further purified by steam distillation. Collected fractions were adjusted to pH 7.0 using NaOH. Pure MGO concentration was indirectly quantified by reaction with H₂O₂ followed by KMnO₄ titration (*60*).

### Cell culture and treatments

HeLa WT cells, HeLa Kyoto EGFP-Lamin A/H2B-mCherry cells (300921, Cytion), and HeLa lamin A/C^KO^ cells (ab261787, Abcam) were routinely cultured at 37 °C and 5% CO₂ in Dulbecco’s Modified Eagle Medium (DMEM) high glucose GlutaMAX (10566016, Gibco) supplemented with 10% fetal bovine serum (FBS) and 1% penicillin/streptomycin. Cells were passaged using Ca^2+^- and Mg^2+^-free Dulbecco’s Phosphate-Buffered Saline (DPBS) (14190144, Gibco) and 0.05% Trypsin-EDTA, except for HeLa Kyoto EGFP-Lamin A/H2B-mCherry cells, which were detached using Accutase solution (400–600 U/mL). Cells used for experiments were maintained between passages 5 and 10.

Unless otherwise specified, cells were seeded the day before any treatment and allowed to adhere overnight before exposing them to the external agent. For MGO exposure experiments, culture medium was replaced with DMEM high glucose without glutamine supplemented with non-essential amino acids (10938025, Gibco) and the indicated concentration of MGO. Cells were incubated under these conditions for 72 h before subsequent experiments, including AFM measurements, confocal microscopy, or nuclear isolation. For histone deacetylase inhibition experiments, cells were incubated in complete culture medium supplemented with 2.5 μM SAHA (HY-10221, MedChemExpress) for 24 h prior to downstream experiments.

### Animal research

#### Diet-induced model

Eight-week-old male C57Bl6/N mice were fed a Western diet (high in fat and sucrose) for 16 weeks (Research Diet D12079B), with monthly monitoring of blood glucose levels and echocardiography. Mice were euthanized and their heart was quickly removed, washed, freeze-clamped and stored at –80 °C.

#### Genetic model

Genetically predisposed obese mice (Lep^ob^ or *ob*/*ob*) carrying spontaneous mutation in leptin were purchased from Charles River (line #606). Heterozygous *ob*/*+* mice were crossed with TEVs-TTN mice, which express a modified version of titin incorporating a Halo ligand binding site and a Tobacco Etch Virus protease cleavage site within its elastic domain (*61*). Descendants were intercrossed to generate homozygous *ob*/*ob* mice and *Lep* WT littermate controls. We used female mice aged 21-26 weeks, euthanized using CO_2_ inhalation. All experimental procedures were performed in accordance with Spanish and European legislation and approved by Comunidad de Madrid ethics committee (PROEX 107.8/23). Within each leptin-genotype group, one animal was heterozygous for the TEVs-TTN allele, whereas the remaining two animals were wild type at the TEVs-TTN locus (WT1 and *ob*/*ob*1: TEVs-TTN^+/-^, heterozygous WT2, WT3, *ob*/*ob*2 and *ob*/*ob*3 TEVs-TTN^-/-^, homozygous). No systematic difference in nuclear mechanical properties or anti-MGO signal was apparent between TEVs-TTN heterozygous and TEVs-TTN WT animals (**Figure S8g-n**).

### Cytotoxicity and oxidative stress assays

#### MTT cytotoxicity assay

Cell viability after MGO exposure was assessed using the Cell Proliferation Kit I 3-(4,5-dimethylthiazol-2-yl)-2,5-diphenyltetrazolium bromide (MTT) (11465007001, Roche). HeLa WT cells were seeded at 0.2 × 10⁶ cells/mL in 12-well plates using DMEM high glucose without glutamine and without phenol red (31053028, Gibco). After overnight adhesion, medium was replaced with 500 μL of MGO-containing medium at final concentrations of 0, 1, 10, 100, or 500 μM. Following 72 h incubation, 100 μL MTT reagent solution (5 mg/mL in DPBS) was added to each well and incubated for 4 h at 37 °C. Formazan crystals were dissolved using solvent solution containing 4 mM HCl and 0.1% NP40 in isopropanol. Plates were shaken at 120 rpm for 30 min and thoroughly mixed to ensure complete dissolution of the formazan crystals. Absorbance was measured at 570 nm using 690 nm as reference wavelength. As a positive control for cytotoxicity, cells were treated with 0.5 ng/mL TNFα for 24 h before MTT measurement. Negative controls included samples without MTT and wells without cells.

#### Intracellular ROS quantification

Reactive oxygen species (ROS) production was assessed using the 2’,7’-dichlorofluorescin diacetate (DCFDA)/ 2’,7’-dichlorodihydrofluorescein diacetate (H2DCFDA) Cellular ROS Assay Kit (ab113851, Abcam). HeLa WT cells were seeded at 0.35 × 10⁶ cells/mL in 48-well plates and allowed to adhere overnight. Cells were then incubated for 72 h with MGO concentrations of 0, 1, 10, 100, or 500 μM. Cells were incubated with 10 μM H2DCFDA in DPBS for 10-20 min followed by two DPBS washes. Positive control cells were treated with 500 µM TBHP for 45 min before staining. Negative controls included samples without H2DCFDA and wells without cells. Cells were fixed using 4% paraformaldehyde (PFA) and counterstained with Hoechst 33342 (1:1000, Invitrogen). Images were acquired using an Axio Observer Z1 epifluorescence microscope. 2’,7’-dichlorofluorescein (DCF) fluorescence intensity was quantified using ImageJ software after background subtraction and autofluorescence correction.

### Nuclear isolation and sample preparation

#### Nuclear isolation from cultured cells

For nuclear isolation, 2.8 × 10⁶ cells were seeded in 100 mm culture dishes (353003, Corning) and cultured for 24 h prior to extraction. In the case of chronic glycation experiments, cells were cultured for 72 h in medium containing 500 µM MGO. For SAHA-treated samples, cells were incubated with 2.5 μM SAHA for 24 h. Paired untreated controls were cultured for the same amount of time.

Prior to extraction, cells were incubated with 1 μM Lat A (L5163, Sigma-Aldrich) for 45 min at 37°C to facilitate release of intact nuclei through actin cytoskeleton disruption. Nuclei were isolated using the Nuclei EZ Prep kit (NUC101, Sigma-Aldrich). Cells were lysed twice using EZ Lysis Buffer and nuclei were collected by centrifugation (1000 × g, 5 min, 4 °C). Isolated nuclei were stored at −80 °C in EZ Storage Buffer for a maximum of two months. Thawed aliquots were not refrozen.

#### Nuclear isolation from tissue

Nuclei from heart tissue were isolated using the Nuclei PURE Prep kit (NUC201, Sigma-Aldrich). Approximately 20 mg tissue was minced and mechanically homogenized using a Dounce homogenizer pestle B in Nuclei PURE Lysis Buffer. Nuclei were collected by centrifugation (1000 × g, 5 min, 4 °C) and further purified over a 2 M sucrose cushion by centrifugation (4100 × g, 50 min, 4 °C, swinging rotor).

Purified nuclei were resuspended in Nuclei PURE Storage Buffer, centrifuged again (1000 × g, 5 min, 4 °C), and stored at −80 °C until use. Thawed aliquots were not refrozen.

### Preparation of isolated nuclei for AFM and microscopy

For AFM experiments, isolated nuclei were immobilized on poly-L-lysine (PLL)-coated (molecular weight 70.000-150.000 Da, P6282, Sigma) glass-bottom dishes (GWST-3522, Wilco). Dishes were coated with 1% PLL in DPBS for 20 min at 37 °C, washed with DPBS, and dried under N₂ flow. Approximately 1 × 10⁶ nuclei in 1 mL DPBS were added to coated dishes and allowed to adhere for 30 min. Unattached nuclei were removed by two DPBS washes.

For *ex-cellulo* glycation experiments, nuclei were incubated for 6 h in either 50 mM MGO in DPBS or DPBS alone. When high-phosphate buffer was used, nuclei were incubated under the same conditions in a buffer containing NaCl (8 mg/mL), KCl (0.2 mg/mL), HPO_4_²⁻ (3.42 mg/mL), and H₂PO_4_⁻ (0.6 mg/mL), with an osmolarity of 360 mOsm/kg and a pH of 7. After incubation, nuclei were washed three times with DPBS prior to AFM measurements or fixation for immunofluorescence.

### Preparation of living cells for AFM and microscopy

For AFM and confocal microscopy experiments, 28,000 corresponding cells (HeLa WT or HeLa Kyoto) were seeded onto 35 mm polymer-bottom dishes (80136, Ibidi). After overnight adhesion, medium was replaced with the respective treatment conditions, and cells were incubated for the required duration before AFM measurements or fixation for immunofluorescence.

### YAP mechanosensing experiments

HeLa WT cells were seeded onto Ibidi dishes as described above and treated with 500 µM MGO for 72 h. Cells were then detached using 0.05% Trypsin-EDTA and resuspended in glutamine-free DMEM. Cells in suspension were replated onto commercially available polyacrylamide gels with elastic moduli of 0.1 kPa or 50 kPa (SV3520-EC-0.1 and SV3520-EC-50, Cell Guidance Systems) and allowed to adhere overnight prior to fixation and YAP immunostaining.

### Atomic force microscopy

AFM measurements were performed using a NanoWizard IV AFM system (Bruker-JPK) coupled to an inverted Axio Observer A1 microscope (Carl Zeiss). For isolated nuclei, pre-calibrated cantilevers with spherical probes of 1 μm radius and spring constant of 0.03 N/m (MLCT-SPH-1UM-DC-D, Bruker) were used. For living cells, pre-calibrated cantilevers with spherical probes of 5 μm radius and approximately 0.3 N/m spring constant (MLCT-SPH-5UM-F, Bruker) were used. Cantilever sensitivity was determined from the thermal noise spectrum (*62, 63*).

Force–distance curves were acquired in contact mode following optical positioning of the cantilever above the nucleus or nuclear region of the cell. Measurements were performed at a constant approach/retract speed of 5 μm/s and a sampling frequency of 2048 Hz. Indentation depths were adjusted to approximately 1.5 μm for isolated nuclei, corresponding to setpoints between 100 pN and 1 nN, and 2.5 μm for intact cells, corresponding to setpoints between 15 and 30 nN. Three consecutive force curves were recorded per sample with approximately 30 s waiting time between measurements to allow mechanical recovery. For experiments assessing the contribution of the actin cytoskeleton, intact cells were first measured under basal conditions and subsequently incubated with 1 μM Lat A for 20 min prior to a second measurement set. For all the experiments, a paired and a treated sample was measured on the same day. Cells were maintained at 37 °C, 5% CO_2_ and immersed in culture medium throughout the course of the experiment. For all mechanical measurements, at least 20 nuclei or cells were measured at each replicate.

Simultaneous AFM and confocal microscopy experiments were performed using a NanoWizard V AFM system (Bruker-JPK) coupled to a LSM900 confocal microscope (Carl Zeiss) equipped with a C-Apochromat 63×/1.20 W Corr M27 objective.

AFM indentation was performed using the same cantilevers and approach parameters described above. During these experiments, force was maintained constant for approximately 60 s while confocal z-stacks were acquired. Because of the force-clamp protocol, these curves were not used for viscoelastic quantification. EGFP-lamin A/C and H2B-mCherry fluorescence were excited using 488 nm and 561 nm lasers, respectively, both at 0.2% laser power. Pinhole size was set to 52 μm. Pixel dwell time was 4.12 μs, with XY pixel size of 0.05 μm and z-step size of 0.5 μm.

### Immunofluorescence staining

Isolated nuclei were fixed using 1.6% PFA (Image-iT Fixative Solution, Thermo Fisher) for 3 min at room temperature and permeabilized using 70% ethanol for 20 min at 4 °C. Cells were fixed using 4% PFA for 10 min at room temperature and permeabilized using 0.2% Triton X-100 in 1% BSA-DPBS for 5 min.

Following permeabilization, samples were washed three times with DPBS and blocked for 1 h at room temperature using SB-buffer containing 10% inactivated FBS and 13.6 g/L BSA in DPBS. Samples were incubated overnight at 4 °C with the corresponding primary antibody. After washing with SB-buffer, samples were incubated with secondary antibodies for 1 h at room temperature when required. Samples were finally washed with DPBS and counterstained using Hoechst 33342 (1:1000). The following antibodies were used in this study: Mouse Methylglyoxal Monoclonal Antibody (1:200, STA-011, Cell Biolabs), lamin A/C Alexa Fluor 488 (1:500, sc-376248, Santa Cruz Biotechnology), Mouse anti-YAP 63.7 (1:100, sc-101199, Santa Cruz Biotechnology), MG-H1 Monoclonal Antibody 1H7G5 (1:50, MA5-49069, Thermo Fisher Scientific), and Goat anti-Mouse Alexa Fluor Plus 647 secondary antibody (1:2000, A32728, Thermo Fisher).

### Confocal microscopy acquisition

Confocal microscopy experiments were performed using a LSM980 Airyscan 2 (Carl Zeiss) equipped with either a C-Apochromat 63×/1.20 W Corr M27 objective or a C-Apochromat 40×/1.2 W Corr M27 objective.

Unless stated otherwise, the central nuclear plane was selected for imaging. Fluorophores were sequentially excited using independent tracks to minimize cross-excitation. Hoechst, AF488, and AF647 fluorophores were excited using 405 nm, 488 nm, and 639 nm lasers, respectively. Laser power was adjusted according to imaging conditions on each experimental day. Therefore, all measurements were performed using paired control and treated samples acquired with identical microscope settings. At least 20 nuclei or cells from at least five different regions of the sample were measured in each experiment. Quantifications were normalized to the corresponding paired control condition. YAP quantification experiments were performed using z-stack acquisition with 1 μm z-resolution.

### Data analysis

#### Power-Law-Rheology (PLR) viscoelastic characterization

Force-distance curves from AFM experiments were analyzed to extract viscoelastic parameters using an in-house Python code. Raw curves were submitted to contact point determination, approach baseline correction and tip displacement rectification. The final processed curve was fitted to a single power-law rheology model characterized by two parameters, scaling modulus (*E*_0_) and fluidity (*γ*). The model accounts for the geometry of the tip and the thickness of the specimen and its analytical expression to determine force as a function of time is as follows (*27*):

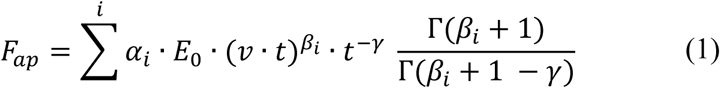

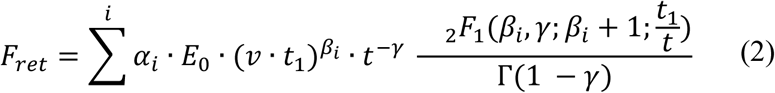

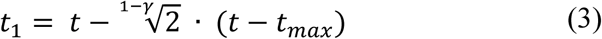

Where *F*_ap_ and *F*_ret_ denote the approach and retract force–distance curves, respectively. Other parameters include *v*, which denotes the indentation velocity, *t* is the time of measurement and *t*_max_ the time point at which the approach phase ends and the retract phase begins. *Γ* refers to the gamma function, and *_2_F_1_* to the hypergeometric function. The coefficients *α_i_* and *β_i_* account for bottom-effect corrections and depend on both the sample height (*h*) and the geometry of the AFM tip. For spherical tips of radius *R*, the coefficients are the following (*26*):

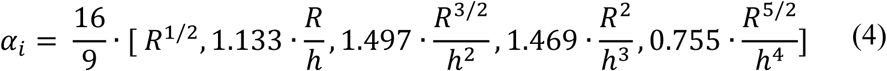

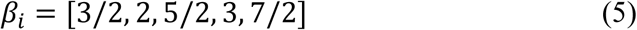

Sample height (*h*) was determined statistically from confocal microscopy z-stacks for each sample. Every nucleus/cell was measured three times and the mean value of *E*_0_ and *γ* after fitting was extracted to represent its viscoelastic properties.

#### Fluorescence intensity quantification

All fluorescence intensity quantifications performed in this study were analyzed using the open-source Fiji software platform (*64*). Unless otherwise specified, nuclear segmentation was performed from the Hoechst channel using the default thresholding algorithm implemented in Fiji. The resulting binary mask was subsequently processed using the Find Particles function to identify individual nuclei. Segmentation outputs were visually inspected to exclude potential non-nuclear particles or segmentation artifacts.

For each image, three independent background regions were selected, and their mean fluorescence intensity values were measured. The average intensity of these regions was defined as the background signal. Nuclear fluorescence intensities were background-corrected by normalization to this mean background value.

To minimize variability arising from day-to-day experimental or imaging conditions, control and treated samples were always acquired in parallel on the same day using identical microscope settings. Following background normalization, the median fluorescence intensity value of the paired control condition was calculated and used as an internal normalization reference. Fluorescence intensities from all nuclei within the corresponding experimental pair were then normalized to this control median value. Final fluorescence measurements are therefore reported as fold change relative to the paired control condition.

For YAP localization experiments, confocal z-stacks were first converted into maximum intensity projections prior to image analysis. Nuclear segmentation was performed from the Hoechst channel using Fiji, while cellular contours were manually delineated using both YAP fluorescence and brightfield images as references. The cytoplasmic region was subsequently defined as the cellular area excluding the nuclear mask. Mean fluorescence intensities within the nuclear and cytoplasmic compartments were measured for each individual cell, and YAP nuclear-to-cytoplasmic (N/C) ratios were calculated accordingly.

#### Nuclear condensate displacement analysis

For nuclear condensate displacement analysis, confocal z-stacks acquired during simultaneous AFM-confocal experiments were analyzed using the Histone 2B fluorescence channel. Raw image stacks were first processed using the Cellpose3 deblurring algorithm with a diameter parameter of 15 px. Enhanced image stacks were subsequently imported into Fiji and organized as hyperstacks containing both z-position and indentation state dimensions (non-indented versus indented states). Indented state image was used to determine the lowest z-plane in which the AFM tip was visibly deforming the nucleus. This position was manually defined and stored in the Fiji ROI Manager. Individual nuclear condensates were then identified within the indented nucleus. For each such condensate, the z-plane displaying the sharpest contour definition was selected manually. In ambiguous cases, orthogonal XZ and YZ projections were used to improve localization accuracy. Then, regions of interest corresponding to each nuclear condensate were manually drawn in the indented state and their XYZ centroid coordinates were measured. The same procedure was repeated for all condensates present within the analyzed nucleus.

The corresponding non-indented state within the hyperstack was subsequently analyzed. Previously defined ROIs were used as spatial references to identify the corresponding condensate positions prior to indentation. Condensates were tracked manually based on their characteristic morphology and relative spatial organization within the nucleus. ROI positions were updated accordingly, and XYZ centroid coordinates were measured again. Condensate displacement was calculated as the three-dimensional Euclidean displacement between the indented and non-indented centroid positions. Additionally, the spatial distance between the AFM tip indentation site and the final condensate position was calculated to assess the spatial propagation range of the applied mechanical perturbation.

#### Nuclear morphology and curvature quantification

To quantify nuclear morphological parameters XZ and YZ orthogonal projections were generated in Fiji from confocal z-stacks using the combined lamin A/C and Histone 2B fluorescence channels. For morphological analysis, projections were generated through the center of the nucleus, defined from the central point of its middle XY plane. For the analysis of nuclear curvature during AFM indentation, the projections were instead centered on the visible AFM indentation site.

Projection images were then analyzed using a custom-written Python script. Images were thresholded to segment the nucleus and extract its contour. Nuclear height was calculated as the vertical distance between the highest and lowest points of the extracted contour, whereas nuclear width was calculated as the horizontal distance between its leftmost and rightmost points.

Apical curvature was determined from the upper nuclear contour. The highest point of the contour was first identified and used as the center of the fitting region. A region comprising 30% of the contour points surrounding this position (±15% on either side) was selected and fitted with a second-order polynomial, *ax*^2^ + *bx* + *c*. The apical curvature was then calculated as *2a*.

For the analysis of the local radius of curvature during AFM indentation, three points along the indentation profile were manually selected by the user: the point of maximum indentation and two lateral points defining the local nuclear contour on either side of the indentation site. These three points were fitted with a second-order polynomial fit as before. The local radius of curvature at the indentation apex was subsequently calculated from the fitted parabola as: 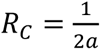.

All morphological and curvature quantifications were performed independently on the XZ and YZ projections, and the mean of the two values was used as the representative apical curvature for each nucleus.

#### Mechanical solidity index (MSI)

Mechanical solidity index (MSI) was obtained from the standardized differences in *E*_0_ and *γ* between basal and altered conditions, corresponding to −MGO vs +MGO, chow vs WD, and WT vs *ob/ob*, respectively. For each mechanical parameter, the difference between conditions was first standardized relative to the variability of the basal condition:

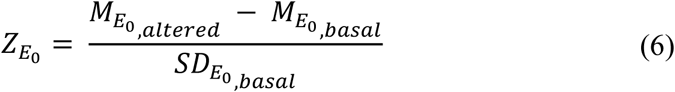

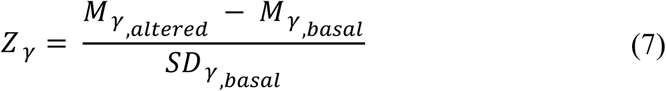

where *M* is the median of the distribution and *SD* is the standard deviation of the basal sample distribution. These differences were combined to calculate the MSI as follows:

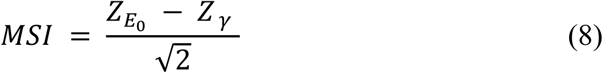

#### Statistical analysis

All statistical analyses were performed using two-tailed Mann–Whitney U test as the primary statistical test for comparisons between two independent groups, as normality could not be consistently assumed across the different experimental datasets. Statistical significance was defined as p < 0.05. In all figures, statistical significance is indicated as follows: p < 0.05 (*), p < 0.01 (**), and p < 0.001 (***).

## Acknowledgments

We thank all members of the nBIO lab and collaborators who contributed to this work.

## Funding

‘La Caixa’ Foundation (LCF/BQ/PI22/11910029, A.C.D).

European Union’s Horizon Europe research and innovation programme, Marie Skłodowska-Curie grant (No. 101065552, DILEMMA, A.C.D).

Research Associate of the F.R.S.-FNRS (A.C.D).

Fonds de la Recherche Scientifique – (FNRS-MIS F402124F, A.C.D).

Research Department of the Communauté française de Belgique (Concerted Research Action MEDACOM, A.C.D).

Ministerio de Ciencia, Innovación y Universidades (MICIU, MICIU/AEI, grant PID2022-140352NA-I00, A.C.D).

Fonds de la Recherche Scientifique (FSR24, A.S.T).

Fonds de la Recherche Scientifique (FSR26, E.M).

Fund for Research Training in Industry and Agriculture (FRIA, FC57485).

Comunitat Autonòma de les Illes Balears, Ajut per a projectes de recerca científica i tecnológica (PRD2025_101, M.A).

Instituto de Salud Carlos III (ISCIII) (project PI25/00843, M.A.).

Ministerio de Ciencia, Innovación y Universidades (MICIU, MICIU/AEI/10.13039/501100011033, European Regional Development Fund, ERDF (PID2023-147683NB-I00, J.A.C).

Instituto de Salud Carlos III (ISCIII, J.A.C).

MICIU (J.A.C).

Pro CNIC Foundation (J.A.C).

Severo Ochoa Center of Excellence (grant CEX2020-001041-S, J.A.C).

Spanish Ministry of Science, Innovation and Universities, José Castillejo Mobility Grant (CAS24/00194, O.M.B.A).

Postdoctoral Researcher, Fonds Recherche Scientifique (N.F).

## Author contributions

Conceptualization: ACD, MAH

Methodology: ACD, MAH

Investigation: MAH, ACD, AD, AST, EM, GJ, RSR, NF, MA, OMBA

Validation: MAH, ACD, AD, AST, EM, GJ, RSR, NF, MA, OMBA

Formal analysis: MAH, ACD, AD, JRR, AST, EM, GJ

Visualization: MAH, ACD

Data curation: MAH, ACD

Software: MAH, JRR

Resources: ACD, JAC, LB, MA, OMBA

Writing – original draft: ACD, MAH

Writing – review & editing: All authors

Supervision: ACD, JAC, LB

Funding acquisition: ACD, JAC, LB, MA, OMBA

Project administration: ACD

## Data, code, and materials availability

The authors declare that the data supporting the findings of this study are available within the paper and its Supplementary Information files. Should any raw data files be needed in another format they are available from the corresponding authors upon reasonable request.

