## Supplementary material for "Methylglyoxal-derived glycation tunes nuclear viscoelasticity"

Mar Alcaraz-Hurtado *et al.*

**This PDF file includes:**

Figs. S1 to S9

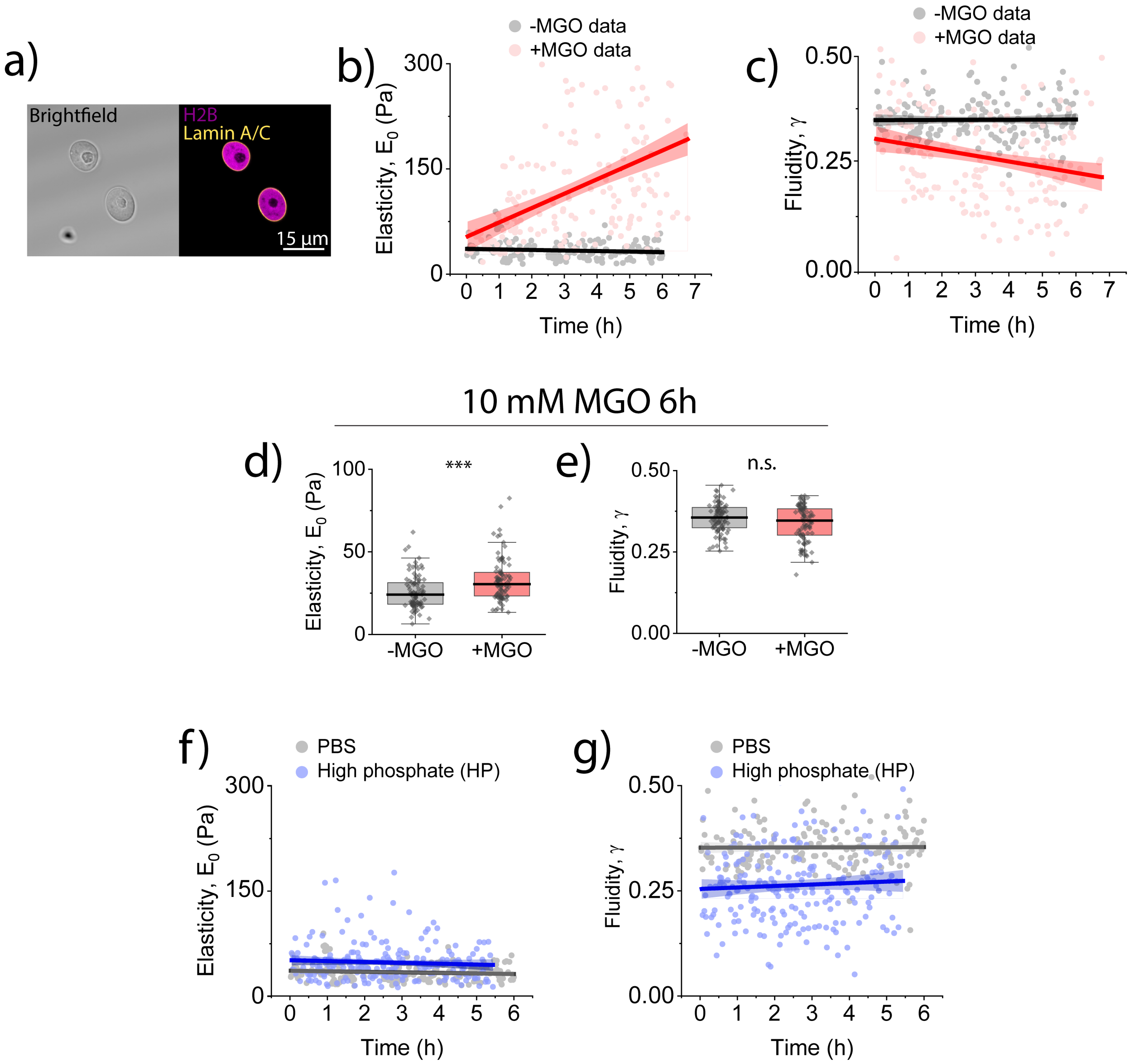


**Fig. S1. MGO-derived glycation remodels nuclear viscoelasticity in a time- and concentration-dependent manner. (a)** Representative brightfield and confocal images of isolated nuclei from HeLa Kyoto stably expressing lamin A/C-EGFP and H2B-mCherry **(b)** Time-dependent quantification of power-law rheology parameters: scaling modulus $E_0$ and **(c)** fluidity coefficient $\gamma$ for untreated (−MGO) and MGO-treated (+MGO, 50 mM) isolated nuclei. Individual points represent the mean value of three consecutive measurements on the same nucleus. Data were pooled from three independent measurements (N(-MGO)=206 nuclei, N(+MGO)=171 nuclei). Gray and red lines represent the linear trend of the data over time, with their corresponding 95% confidence intervals (CI). **(d)** Quantification of the scaling modulus $E_0$ and **(e)** fluidity coefficient $\gamma$ for isolated nuclei untreated (-MGO) or treated (+MGO) with MGO 10 mM for 6h. Each dot represents the mean of three consecutive measurements per nucleus. Data were pooled from three independent measurements (N(-MGO)=90 nuclei, N(+MGO)=89 nuclei). Boxes indicate 25th–75th percentiles, whiskers indicate outlier boundary and the median of the distribution is represented with a thicker horizontal line. **(f)** Time-dependent quantification of power-law rheology parameters: scaling modulus $E_0$ and **(g)** fluidity coefficient $\gamma$ for nuclei in PBS (gray) and in high phosphate buffer (blue), as a hyperosmotic/high-phosphate control. Individual points represent the mean value of three consecutive measurements on the same nucleus. Data were pooled from three independent measurements (N(-MGO)=206 nuclei, N(+MGO)=213 nuclei). Gray and blue lines represent the linear trend of the data over time, with their corresponding 95% confidence intervals (CI). Statistical significance was assessed using Mann–Whitney tests; *$p < 0.05$, ***$p < 0.001$; n.s., not significant.

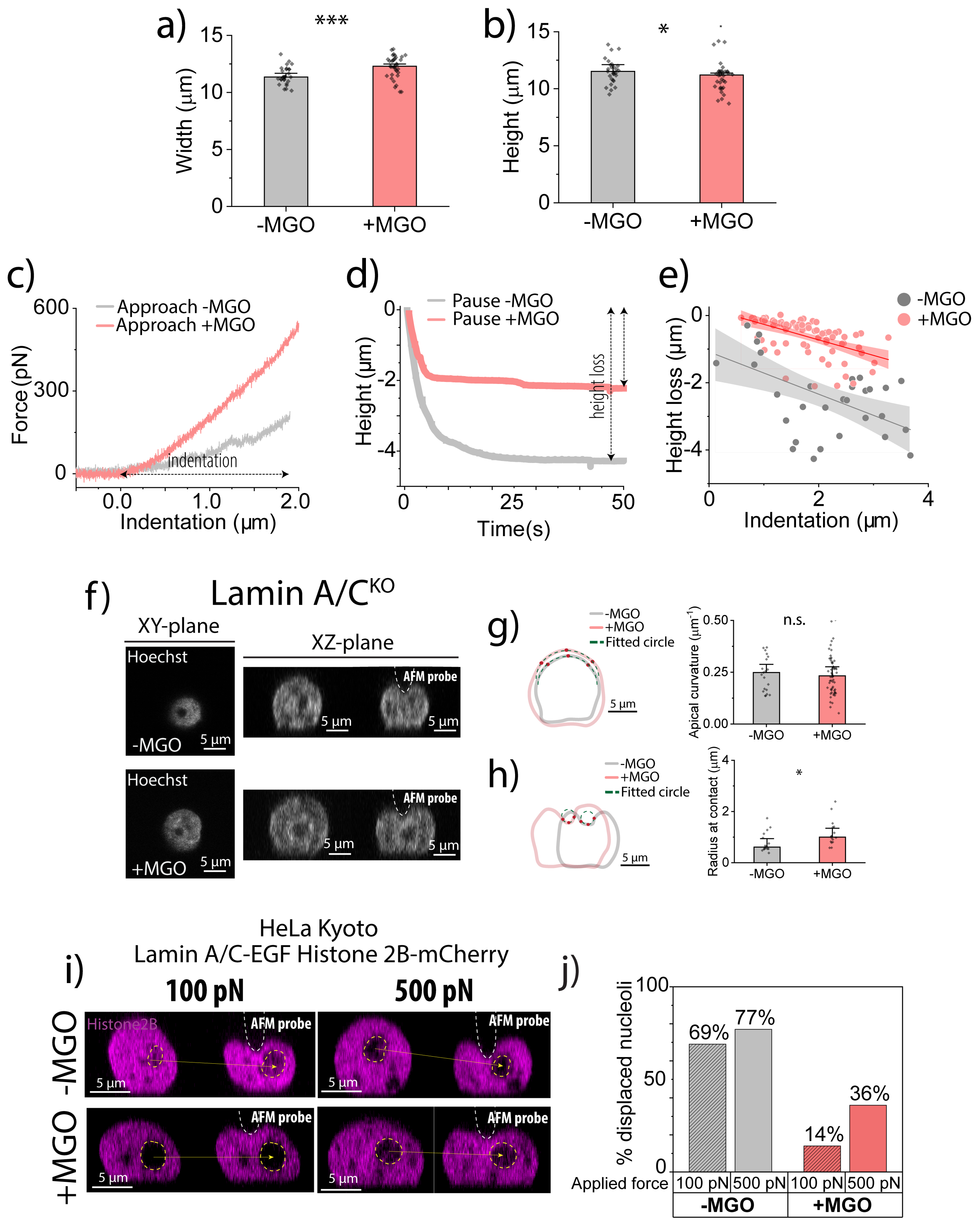


**Fig. S2. MGO-treated nuclei resist local deformation and show reduced intranuclear deformation propagation. (a)** Width and **(b)** height quantification of HK-LAC-H2B nuclei untreated (-MGO) or treated (+MGO) with MGO 50 mM during 6h, before indentation. Each point represents the mean width or height obtained from XZ and YZ projection of one nucleus. Bars show median ± SEM and data were pooled from at least three independent experiments (N(-MGO)=29 nuclei, N(+MGO)=35 nuclei). **(c)** Representative force-indentation approach segment of untreated (-MGO) and treated (+MGO) HK-LAC-H2B nuclei. Similar indentations (around 2 μm) require the application of higher forces in +MGO nuclei. **(d)** Representative time-evolution of height during pause segment of untreated (-MGO) and treated (+MGO) HK-LAC-H2B nuclei. These pause segments belong to the same curves as the approach segments represented in **(c)**. Similar indentations produce reduced height loss in MGO treated nucleus. **(e)** Quantification of height loss as a function of indentation depth in untreated (-MGO) and treated (+MGO) nuclei. Each dot represents the measurement of the indentation and height loss in a single curve. Gray and red lines represent the linear trend of the data over time, with their corresponding 95% confidence intervals (CI). Data were pooled from three independent experiments (N(-MGO)=24 nuclei, N(+MGO)=77 nuclei). **(f)** Representative confocal z-stacks of untreated (−MGO) and treated (+MGO) isolated HeLa lamin A/$C^{KO}$ nuclei during AFM indentation. Left: XY plane at the indentation site; right: XZ projections before and after indentation. Hoechst signal is shown in white. Dashed white line indicates the AFM probe. **(g)** Apical curvature analysis before indentation of nuclei shown in **(f),** with nuclear contour shown in gray (-MGO) and red (+MGO), fitting limits marked as red dots, and result fitted circle in green. Each point represents the mean curvature from XZ and YZ projections of one nucleus; bars indicate median ± SEM. Data are representative of at least two independent experiments (N(-MGO)=16 nuclei, N(+MGO)=16 nuclei). **(h)** Radius at contact analysis upon indentation of nuclei shown in **(f),** with nuclear contour shown in gray (-MGO) and red (+MGO), fitting limits marked as red dots, and result fitted circle in green. Each point represents the mean radius obtained from XZ and YZ projection of one nucleus. Bars show median ± SEM and data were pooled from at least two independent experiments(N(-MGO)=21 nuclei, N(+MGO)=60 nuclei). **(i)** Representative confocal images of HK-LAC nuclei during 100 and 500 pN indentation. Histone 2B is shown in magenta; dashed yellow outlines a nuclear condensate before and during force application, with displacement indicated by arrows. **(j)** Percentage of nuclear condensates displaced at 100 pN and 500 pN in −MGO and +MGO conditions. Data pooled from three independent paired experiments (N(-MGO, 100 pN)=22 nuclei, N(-MGO, 500 pN)=10 nuclei, N(+MGO, 100 pN)=10 nuclei, N(+MGO, 500 pN)=16 nuclei)). Statistical analysis: Mann–Whitney test; $^{*}p < 0.05$, $^{**}p < 0.01$, $^{***}p < 0.001$, n.s. = not significant.

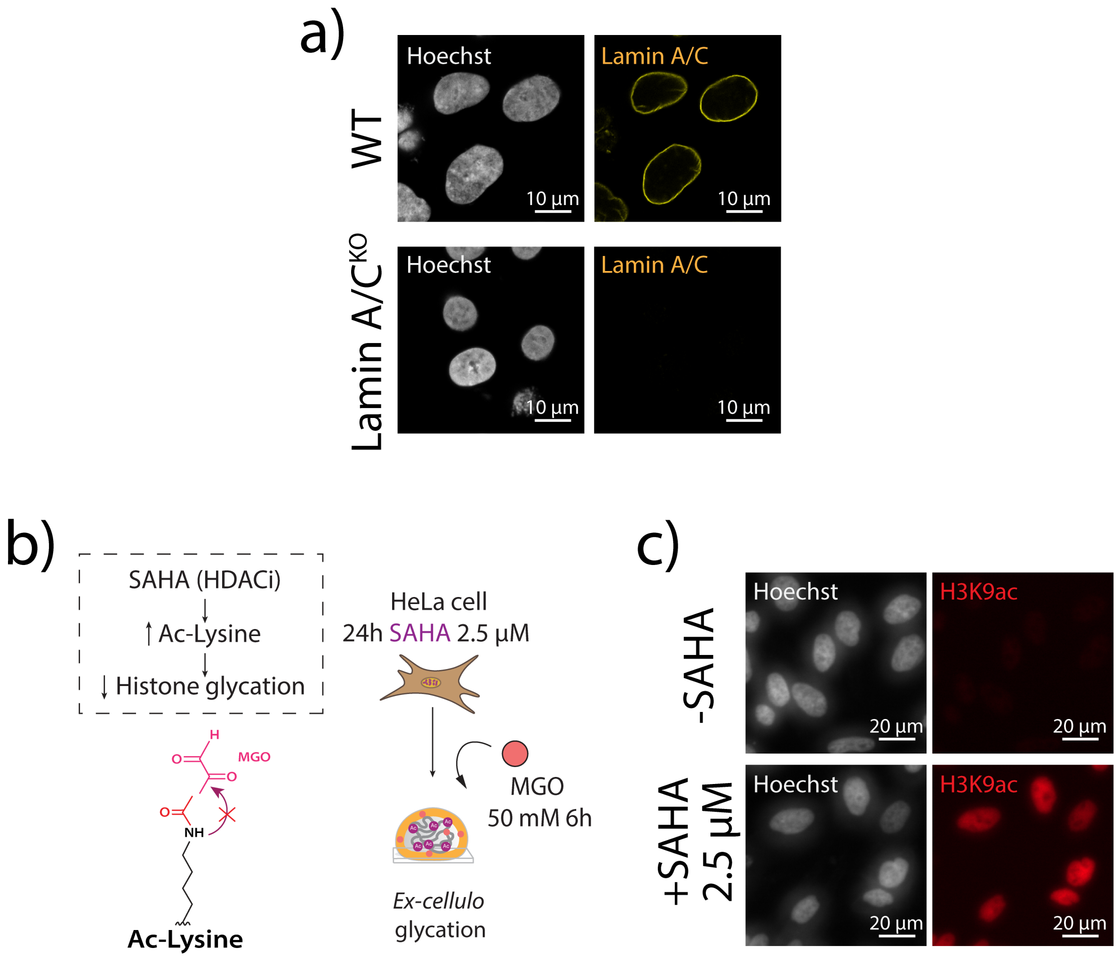


**Fig. S3. Experimental strategies used to probe the relative contributions of nuclear lamina and chromatin. (a)** Representative confocal microscopy images of WT and Lamin A/C$^{KO}$ isolated nuclei immunostained for lamin A/C. **(b)** Schematic of the strategy used to modulate chromatin state and reduce histone lysine availability for MGO-derived glycation. Histone deacetylase inhibitors, such as SAHA, efficiently increase the levels of acetylated lysine in histones, which can reduce lysine availability for MGO-derived glycation. Cells were treated with SAHA 2.5 µM for 24h prior to nuclear extraction and *ex-cellulo* glycation with MGO 50 mM. **(c)** Representative epifluorescence microscopy images of cells treated with SAHA and immunostained for histone H3 lysine 9 acetylation (H3K9ac).

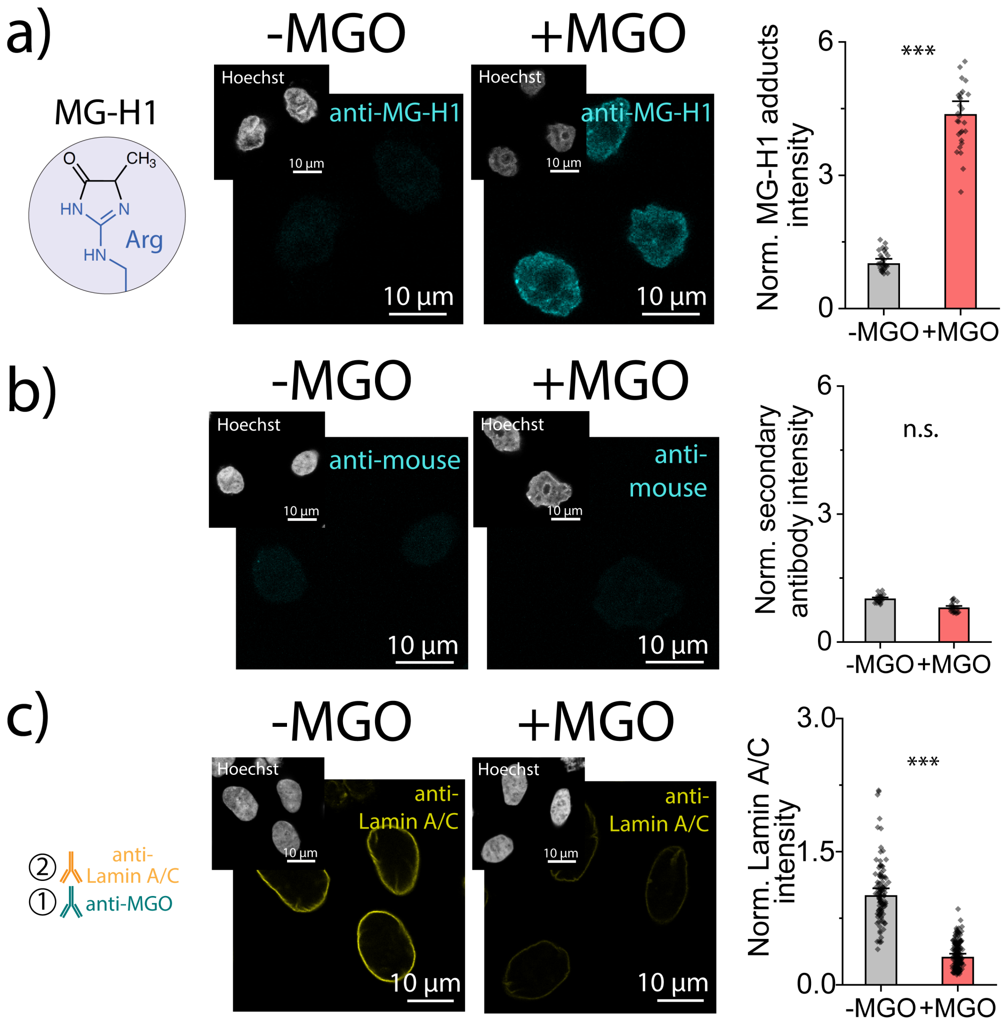


**Fig. S4. MG-H1 and pan-MGO staining validate detection of MGO-derived nuclear adducts.** **(a)** Left: MG-H1 is a specific arginine-derived MGO adduct. Center: representative confocal images of anti-MGH1 signal (blue) with Hoechst inset (white) for −MGO and +MGO conditions. Right: quantification of anti-MG-H1 signal. Each dot represents the mean fluorescence of a single nucleus, background-subtracted and normalized to the median of the −MGO population; bars indicate median ± SEM (N(-MGO)=30 nuclei, N(+MGO)=30 nuclei). **(b)** Representative confocal images of anti-mouse signal (blue) with Hoechst inset (white) for −MGO and +MGO conditions. Right: quantification of anti-mouse signal, analyzed as in **(a)** (N(-MGO)=22 nuclei, N(+MGO)=22 nuclei). **(c)** Representative confocal images of lamin A/C signal (yellow) with Hoechst inset (white) for −MGO and +MGO conditions. Immunofluorescence was performed sequentially staining first MGO-adducts, then lamin A/C. Right: quantification of lamin A/C signal, analyzed as in **(a)** (N(-MGO)=107 nuclei, N(+MGO)=168 nuclei). Statistical analysis: Mann–Whitney test; *$p < 0.05$, **$p < 0.01$, ***$p < 0.001$, n.s. = not significant.

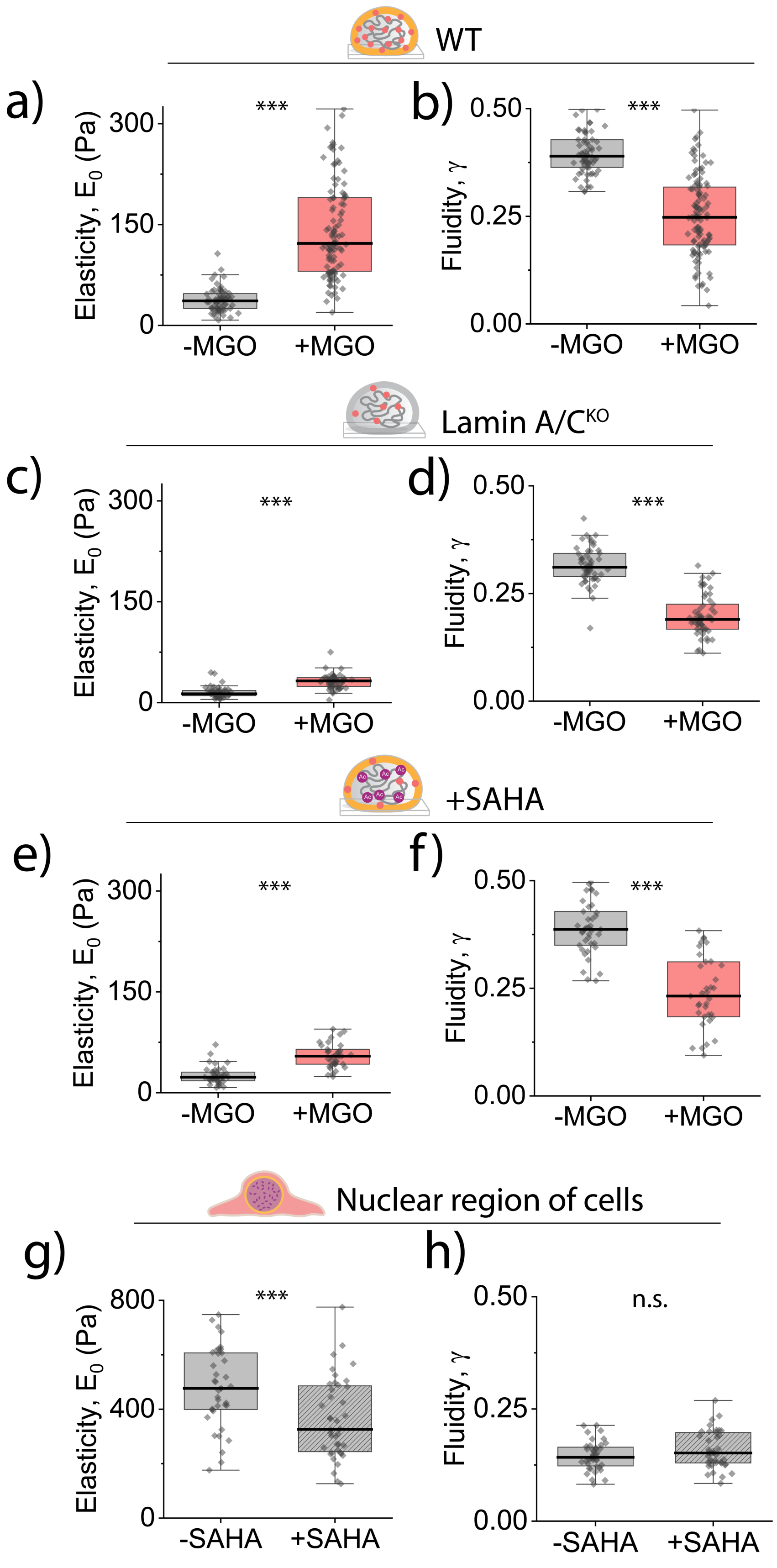
WT
a)
***
300
150
0
Elasticity, $E_0$ (Pa)
-MGO
+MGO
b)
***
0.50
0.25
0.00
Fluidity, $\gamma$
-MGO
+MGO
Lamin A/C$^{KO}$
c)
***
300
150
0
Elasticity, $E_0$ (Pa)
-MGO
+MGO
d)
***
0.50
0.25
0.00
Fluidity, $\gamma$
-MGO
+MGO
+SAHA
e)
***
300
150
0
Elasticity, $E_0$ (Pa)
-MGO
+MGO
f)
***
0.50
0.25
0.00
Fluidity, $\gamma$
-MGO
+MGO
Nuclear region of cells
g)
***
800
400
0
Elasticity, $E_0$ (Pa)
-SAHA
+SAHA
h)
n.s.
0.50
0.25
0.00
Fluidity, $\gamma$
-SAHA
+SAHA

**Fig. S5. Lamin A/C$^{KO}$ and +SAHA nuclei display an attenuated mechanical response to MGO-derived glycation.** **(a)** Quantification of scaling modulus $E_0$ and **(b)** fluidity coefficient $\gamma$ for nuclei isolated from HeLa WT nuclei measured in paired -MGO and +MGO experiments. Each dot represents the mean of three consecutive indentations per nucleus. Data was pooled from three independent measurements (N(-MGO)=64 nuclei, N(+MGO)=109 nuclei). **(c)** Quantification of scaling modulus $E_0$ and **(d)** fluidity coefficient $\gamma$ for nuclei isolated from lamin A/C$^{KO}$ nuclei measured in paired −MGO and +MGO experiments. Each dot represents the mean of three consecutive indentations per nucleus. Data were pooled from three independent measurements (N(-MGO)=52 nuclei, N(+MGO)=55 nuclei). **(e)** Quantification of scaling modulus $E_0$ and **(f)** fluidity coefficient $\gamma$ for nuclei isolated from SAHA treated cells, before (-MGO) and after MGO incubation (+MGO). Each dot represents the mean of three consecutive indentations per nucleus (N(-MGO)=39 nuclei, N(+MGO)=35). **(g)** Quantification of scaling modulus $E_0$ and fluidity coefficient $\gamma$ **(h)** measured in nuclear regions of HeLa cells untreated (-SAHA) or treated (+SAHA) with SAHA 2.5 μM during 24 h. Each dot represents the mean value of three consecutive indentations per cell (N(-SAHA)=40 cells, N(+SAHA)=40 cells). Boxes indicate 25th–75th percentiles, whiskers indicate outlier boundary and the median of the distribution is represented with a thicker horizontal line. Statistical analysis: Mann–Whitney test; *p < 0.05, **p < 0.01, ***p < 0.001, n.s. = not significant.

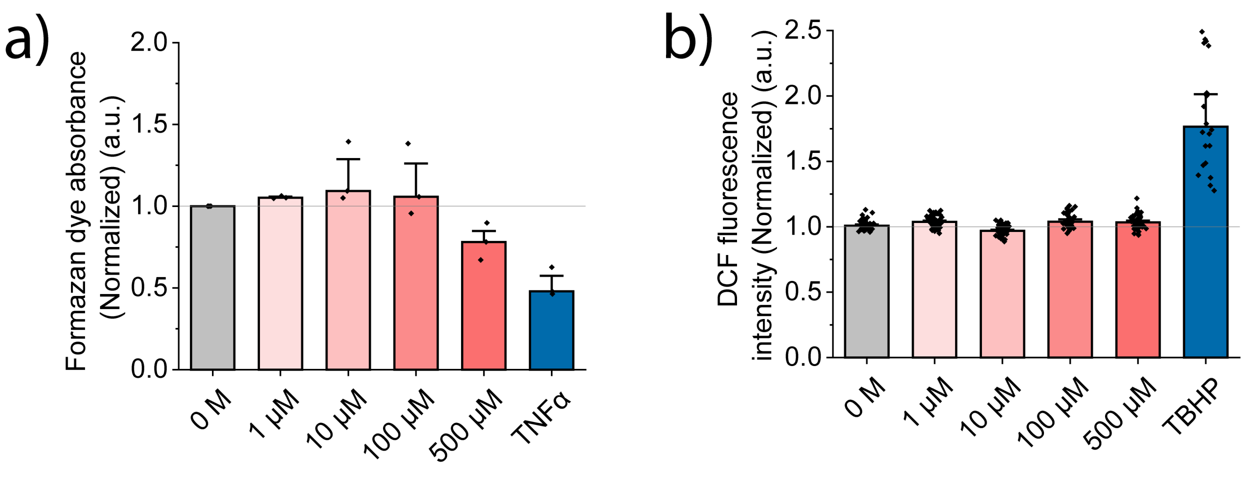


**Fig. S6. Cell viability and ROS generation after chronic MGO exposure for 72 h.** Mitochondrial activity and reactive oxygen species (ROS) were assessed in cells exposed to increasing MGO concentrations (0, 1 µM, 10 µM, 100 µM, 500 µM) over 72 h. **(a)** MTT assay showing mitochondrial activity inferred from solubilized formazan absorbance, with TNF-α as a positive cytotoxicity control. After 72 h exposure, cells were incubated with MTT reagent, formazan crystals were solubilized, and absorbance was measured at 570 nm (three technical readings per well). Each data point represents the mean of the three measurements. Number of biological replicates per condition = 3. **(b)** Intracellular ROS measured with the DCFDA probe; fluorescence of oxidized DCF reflects ROS levels, with TBHP as a positive oxidative stress control. After 72 h exposure, cells were incubated with H2DCFDA, fixed, and stained with Hoechst. Epifluorescence images were acquired, and single-cell fluorescence was quantified using ImageJ, corrected for background and autofluorescence (cells without probe). Each data point represents a single cell. Number of biological replicates per condition = 3. Bars indicate median ± SEM.

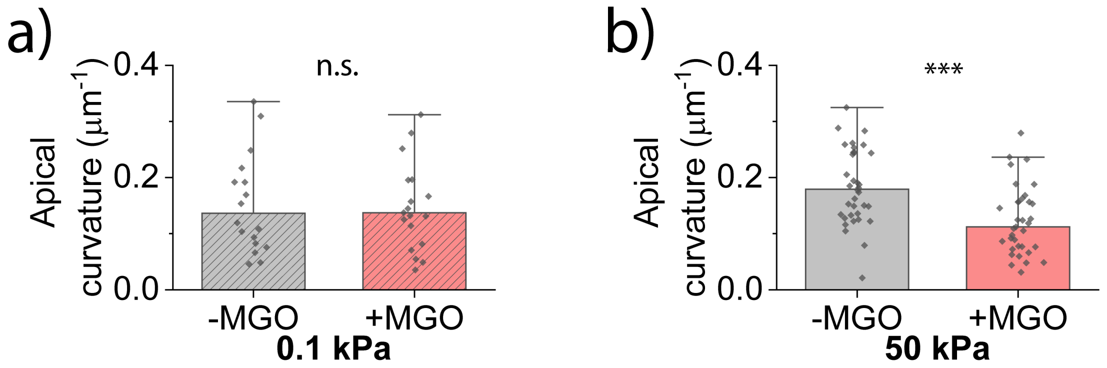


**Fig. S7. Cell geometry after MGO treatment on soft and stiff substrates. (a)** Quantification of cellular apical curvature in -MGO and +MGO cells seeded on soft 0.1 kPa and **(b)** stiff 50 kPa substrates. Each dot represents one cell, and bars indicate median ± SEM (N(-MGO, 0.1 kPa)=30 cells, N(+MGO, 0.1 kPa)=30 cells, N(-MGO, 50 kPa)=39 cells, N(+MGO, 50 kPa)=60 cells). Statistical analysis: Mann–Whitney test; $*p < 0.05$, $**p < 0.01$, $***p < 0.001$, n.s. = not significant.

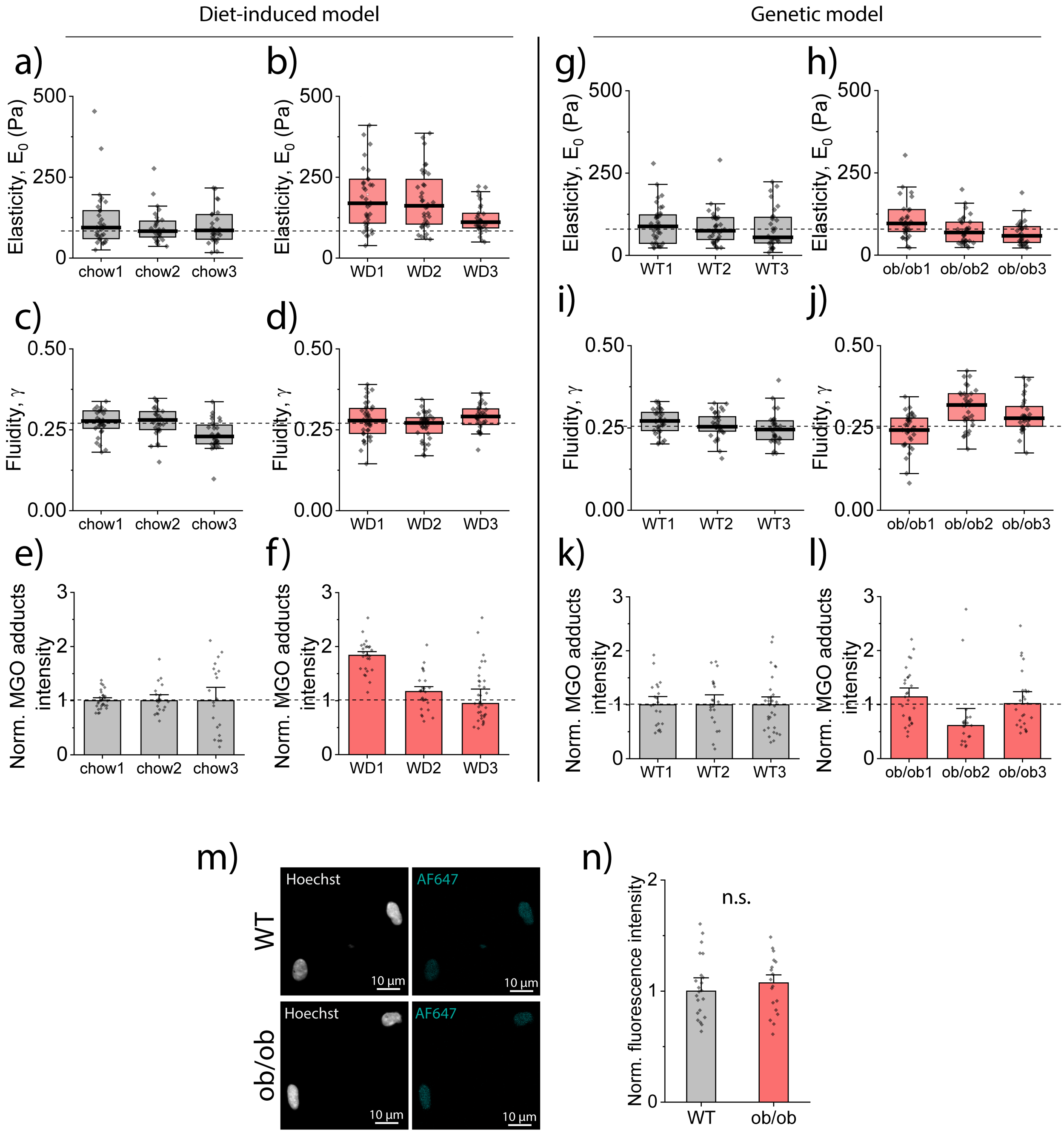


**Fig. S8. Animal-level heterogeneity in cardiac nuclear mechanics and MGO-adduct accumulation. (a-l)** Quantification of scaling modulus $E_0$, fluidity coefficient $\gamma$ and normalized MGO-adducts for nuclei isolated from each of the animals in diet-induced (chow and WD) and genetic model (WT and *ob*/*ob*). WT1 and *ob*/*ob*1 mice are heterozygous with TEVs-TTN, while WT2, WT3, *ob*/*ob*2 and *ob*/*ob*3 were homozygous mice. No mechanical differences are observed between heterozygous and homozygous mice. For mechanical measurement, each dot represents the mean of three consecutive indentations per nucleus. Boxes indicate 25th–75th percentiles, whiskers indicate outlier boundary and the median of the distribution is represented with a thicker horizontal line. For fluorescence quantification, each dot represents the mean value of a single nucleus, background subtracted and normalized to the median of paired controls. Bars indicate median ± SEM. **(m)** Representative confocal images of secondary antibody, anti-mouse signal (blue) with Hoechst inset (white) for control and diabetic conditions. **(n)** Quantification of anti-mouse signal. Each dot represents the mean fluorescence of a single nucleus, background-subtracted and normalized to the median of the matched control population; bars indicate median ± SEM (N(WT)=22 nuclei, N(*ob/ob*)=18 nuclei).

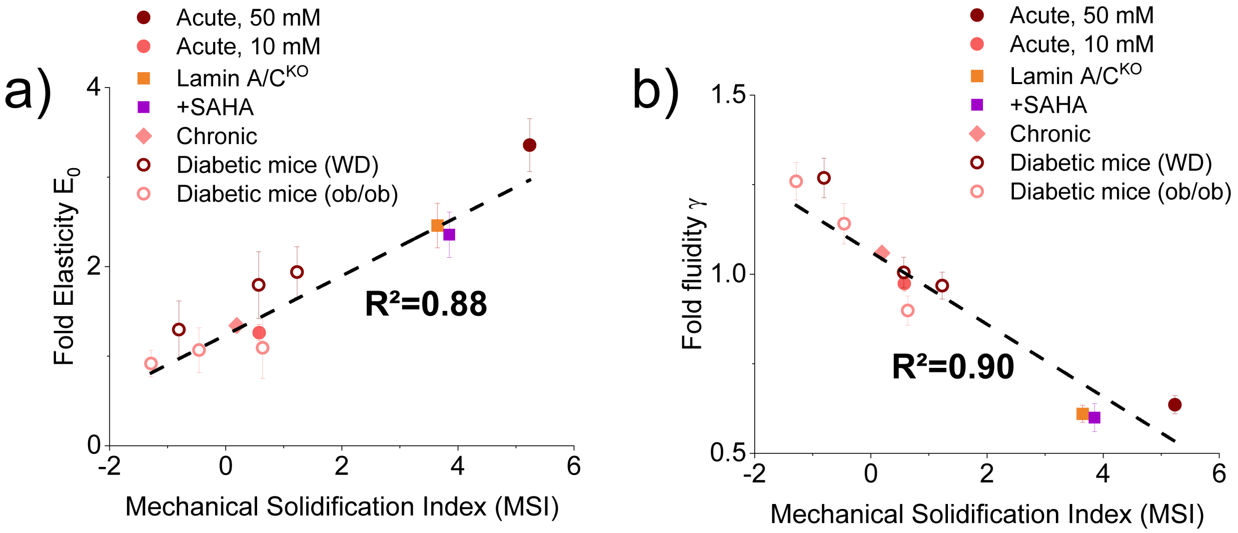


**Fig. S9. MSI captures the combined shift toward increased stiffness and reduced fluidity.** Relationship between the MSI and the individual fold changes in **(a)** scaling modulus and **(b)** fluidity in the carbonyl-stress models analyzed in this study: acute *ex-cellulo* MGO exposure at 50 mM or 10 mM for 6 h, lamin A/C$^{KO}$, SAHA treatment, chronic *in-cellulo* exposure to 500 µM MGO for 72 h and cardiac nuclei from diet-induced and genetic metabolic disease models. MSI was calculated for each transition relative to its paired control as the combined standardized shift toward increased scaling modulus and reduced fluidity, with each term normalized by the control standard deviation. Higher MSI values correspond to a more solid-like nuclear response, whereas values near or below zero indicate little or no shift toward solidification. Error bars indicate SEM of the fold change in scaling modulus or fluidity, calculated by uncertainty propagation from the original SEM values. Dashed lines show linear fits with the corresponding $R^2$.
